# SUZ12–Nucleic Acid Interactions Constrain PRC2 Activity to Maintain Targeted Gene Silencing Essential to Diffuse Midline Glioma

**DOI:** 10.1101/2025.07.18.665585

**Authors:** Tyler J. Reich, Paul A. Clark, Audrey Baguette, Joanna K. Lempiainen, Caterina Russo, Andrew Q. Rashoff, Truman J. Do, Benjamin A. Garcia, Claudia L. Kleinman, Nada Jabado, Zachary S. Morris, Peter W. Lewis

## Abstract

Polycomb Repressive Complex 2 (PRC2) mediates transcriptional silencing through trimethylation of histone H3 at lysine 27 (H3K27me3), an epigenetic modification critical for development and frequently altered in cancer. Pediatric diffuse midline gliomas (DMGs) bearing the histone H3 K27M mutation exhibit global loss of H3K27me3 due to dominant inhibition of PRC2 by the mutant histone. Despite widespread hypomethylation, focal retention of H3K27me3 persists, and tumor cells maintain dependency on residual PRC2 activity for proliferation. The molecular basis underlying this residual enzymatic function and its regulation remain poorly defined. To address this mechanism, we investigated the role of SUZ12, the architectural core of PRC2 that facilitates interactions with accessory subunits. We identified the SUZ12 N-terminal region as a regulatory domain that constrains PRC2 catalytic activity through transient interactions with nucleic acids, thereby limiting non-specific chromatin engagement. Expression of a truncated SUZ12 variant retaining the catalytic VEFS domain, but lacking the nucleic acid-binding regulatory elements, led to widespread H3K27 hypermethylation, displacement of canonical PRC1 complexes, disruption of chromatin architecture, and impaired H3 K27M glioma cell growth *in vitro* and *in vivo*. Biochemical analyses revealed a SUZ12 N-terminal domain that modulates PRC2 activity by promoting non-productive binding to nucleic acids, thus establishing a kinetic equilibrium essential for precise chromatin targeting. These findings redefine Polycomb specificity as a dynamic equilibrium between productive nucleosomal engagement and non-productive nucleic acid interactions, providing critical insights into PRC2 regulation and highlighting potential therapeutic vulnerabilities in PRC2-dependent cancers.

## Introduction

Precise regulation of chromatin structure is essential for establishing and maintaining cellular identity during development and tissue homeostasis. This regulation involves histone-modifying enzymes that deposit, interpret, and remove post-translational modifications (PTMs) on histone tails. Polycomb group (PcG) proteins are critical chromatin regulators that assemble into multimeric complexes to enforce heritable gene silencing^1–3^. Across metazoan development, polycomb repression is indispensable for embryonic patterning, stem cell maintenance, and cell fate decisions and its dysregulation is frequently seen in developmental disorders and cancer^4–7^.

PcG proteins organize primarily into two major complexes, Polycomb Repressive Complex 1 (PRC1) and Polycomb Repressive Complex 2 (PRC2), with distinct yet interconnected functions in chromatin regulation^8^. PRC2 catalyzes mono-, di-, and tri-methylation of histone H3 at lysine 27 (H3K27me1/2/3), while PRC1 chemically modifies chromatin via monoubiquitination of histone H2A at lysine 119 (H2AK119ub)^9^. PRC1 can be further classified into canonical or variant PRC1 complexes (cPRC1 or vPRC1, respectively) by subunit compositions^10^. cPRC1 is defined, in part, by the presence of either PCGF2 or PCGF4 (also known as BMI1) and one of several chromodomain-containing CBX paralogues. Compared to vPRC1, cPRC1 deposits little H2AK119ub in cells and instead is thought to enforce silencing downstream of CBX-H3K27me3 interactions through several described mechanisms, including local chromatin compaction and higher-order genome organization^11–15^.

Core PRC2 is composed of a catalytic subunit EZH1/2 (predominantly EZH2 in dividing cells), along with SUZ12, EED, and the histone-binding protein RBBP4/7^16^. Mammalian PRC2 maintains lineage-specific silencing in a stepwise fashion by first binding CpG island promoters of relevant target genes and depositing high levels of H3K27me3 locally. From initial recruitment sites, PRC2 spreads H3K27me3 via its conserved read-write mechanism, whereby EED-H3K27me3 binding induces conformational rearrangements that increase EZH2 catalysis^17–23^. This trans-subunit allostery represents a key node of PRC2 regulation and its disruption is seen in a range of developmental and malignant diseases^24,25^.

One notable example of this vulnerability occurs in diffuse midline gliomas (DMGs), a class of invariably fatal pediatric brain tumors. A unifying feature of DMGs, seen across molecular and anatomical subtypes, is a unique chromatin profile where H3K27me3 is strictly confined to recruitment sites with concurrent loss across broad spread regions^26^. Approximately 80% of DMGs are driven by monoallelic mutations in histone H3 position 27 (K27M). The resulting K27M oncohistone, accounting for roughly 10% of total H3 protein, drives oncogenic gene expression through potent inhibition of PRC2, resulting in striking reductions in global H3K27me3 levels^22,27,28^. Mechanistically, K27M prevents H3K27me3 spread through its preferential inhibition of allosterically stimulated PRC2^22,29,30^. Of note, most of the remaining H3 wild-type pontine gliomas aberrantly express a soluble protein, EZHIP, that similarly inhibits PRC2 through a conserved mechanism^31,32^.

These insights explain how DMG cells lose H3K27me3 from broad spread regions. By comparison, the other half of the unique chromatin profile in DMGs, recruitment-site H3K27me3, has received little experimental attention. The few studies addressing the pathologic significance of residual H3K27me3 have reached differing conclusions: while some found it to be an attractive dependency, others suggest EZH2 inhibition could hasten tumor growth and worsen patient survival^33–36^. Given this ambiguity and continued lack of clinical options, clarifying the therapeutic relevance of residual PRC2 activity in DMG remains an urgent need. Critically, a more robust understanding of the molecular events responsible for recruitment-site H3K27me3 and associated gene silencing must be established to justify this counterintuitive approach and may inform novel therapeutic avenues.

One barrier to mechanistic insight is the lack of precise molecular descriptions for fundamental PRC2 functions. For example, roles for PRC2-RNA interactions in regulating PRC2 function remain contentious^37^. Similarly, how PRC2 finds its genomic targets remains poorly understood, with only recent insights having defined non-core subunits as essential recruitment factors^38,39^. These accessory subunits comprise PRC2.1 and PRC2.2 subcomplexes in cells^40^. PRC2.1 includes one of three PCL proteins (PHF1, MTF2, or PHF19), which bind CG-rich DNA, and either PALI1 or EPOP^19,41–44^. PRC2.2 contains JARID2 and AEBP2, which enhance binding via crosstalk with PRC1-catalyzed H2AK119ub^45–47^. Accessory subunits compete for binding sites on the N-terminus of core subunit SUZ12, which assembles core members EZH2 and EED through its C-terminal VEFS domain^42,48^. Experiments in embryonic stem cells demonstrated an essential but collective requirement for accessory subunits, where only loss of both subcomplexes (either by combinatorial accessory subunit knockout or expression of a VEFS-only SUZ12 truncation) resulted in complete loss of targeting^38,39,48^. Notably, both genetic approaches produced globally normal methylation levels, suggesting dispensable roles for accessory subunits in maintaining total cellular H3K27me3.

Here we leveraged these insights to comprehensively characterize residual PRC2 activity in K27M DMG. Targeted disruption of PRC2.1/2.2 subcomplexes via SUZ12 mutagenesis revealed an essential role for accessory subunit-mediated recruitment in K27M DMG. Unexpectedly, we identified a novel regulatory element in the SUZ12 N-terminus that restricts cellular H3K27me3 independent of accessory subunits. Expression of VEFS-containing SUZ12 variants lacking this region produced diffusely hypermethylated genomes in all cell types examined. K27M glioma cells were particularly vulnerable to this profile: diffuse H3K27me3 potently dislodged PRC1 from target genes and dominantly disrupted genome architecture with severe cellular consequences. Motivated by the therapeutic implications of this sensitivity, we mapped the domain responsible for restricting diffuse hypermethylation to a zinc finger-containing region in the SUZ12 N-terminus that bound diverse nucleic acid species to inhibit PRC2 catalysis. Collectively, these results (1) unambiguously clarify residual PRC2 activity as a tractable vulnerability in K27M DMG, (2) identify diffuse H3K27me3 as a critical determinant for PRC1 localization irrespective of target-site levels, and (3) describe a novel mechanism of PRC2 catalytic regulation whereby indiscriminate, inhibitory interactions between SUZ12 and nucleic acids constrain pervasive methylation in cells to ensure targeted gene silencing.

## Results

### Residual PRC2 catalytic activity critically sustains growth of K27M-mutant DMG cells

Previous studies evaluating EZH2 inhibitors as therapeutic agents in DMG have yielded inconsistent results^33–36^. To clarify the functional significance of residual PRC2 enzymatic activity in K27M DMG, we conducted comprehensive proliferation assays using patient-derived glioma cell lines expressing either mutant H3 K27M or wild-type histone H3 (**Figure S1A**). Cells were treated with two distinct classes of PRC2 inhibitors: Tazemetostat, a competitive inhibitor targeting the EZH2 catalytic site through the S-adenosylmethionine (SAM) cofactor pocket, and A-395, an inhibitor blocking PRC2 allosteric activation by engaging the aromatic cage of the EED subunit, thereby impairing the PRC2 read-write function.

Notably, proliferation assays were conducted over extended periods (several weeks) to fully deplete preexisting H3K27me3 histone marks through successive cell divisions, given that mammalian cells primarily lose H3K27me3 passively through replicational dilution^49^. Under these prolonged conditions, both Tazemetostat and A-395 caused severe proliferation defects in K27M DMG cell lines, whereas growth rates for wild-type histone H3-expressing gliomas were spared at all inhibitor concentrations tested (**Figure S1B**).

Consistent with previous reports linking PRC2 inactivation to tumor-suppressive pathways^50,51^, treatment with PRC2 inhibitors in K27M mutant DMG cells induced a dose-dependent increase in expression of p16^INK4A^ (hereafter referred to as p16), a tumor suppressor encoded by the cell cycle regulator locus *CDKN2A* (data not shown). Additionally, Tazemetostat exposure resulted in derepression of many genes near PRC2 recruitment sites which were enriched for biological processes related to neurodevelopment and cell cycle regulation (**Figures S1H and S1I**). To directly assess the importance of p16 induction following loss of PRC2 catalysis, we generated clonal p16 knockout cell lines in two distinct K27M mutant DMG models and found that p16 loss largely restored the proliferative defects seen in p16-sufficient cell lines upon PRC2 inhibition (**Figures S1C-S1E**). Together, these findings establish core PRC2 catalytic activity as a requirement for K27M DMG cell growth and p16 as a critical mediator of the cellular processes downstream PRC2 inhibition. However, these experiments do not resolve whether the observed dependency specifically arises from H3K27me3 maintained at focal recruitment sites.

### Accessory PRC2 subunits orchestrate targeted silencing essential for DMG survival

To directly assess the importance of target-site H3K27me3 in DMG, we employed a genetic approach to selectively disrupt PRC2 localization without impairing its intrinsic catalytic function. To this end, we focused on the role of accessory PRC2 subunits in promoting targeted gene silencing essential for DMG cell proliferation. Recent work has shown that non-core/accessory subunits, which define the PRC2.1 and PRC2.2 subcomplexes, are collectively essential for PRC2 localization but dispensable for global methylation levels in cells^38,39^. Accessory subunits compete for binding sites on the core subunit SUZ12, which is thought to primarily act as a scaffolding platform for accessory subunits through its N-terminus and other core subunits (EZH2, EED) through its C-terminal VEFS domain^42,48^.

Leveraging recent biochemical insights, we engineered a panel of SUZ12 mutant constructs to selectively to impair PRC2.1 (2.1mut) or PRC2.2 (2.2mut) assembly, as well as a combined non-core mutant (ncMUT) disrupting both subcomplexes simultaneously^52,53^. Additionally, we included a VEFS truncation control lacking all N-terminal sites needed to dock accessory subunits (**Figure 1A**). We validated each mutant biochemically in co-immunoprecipitation experiments, confirming selective loss of the intended subcomplex while preserving core PRC2 assembly and catalytic competence (**Figures 1B, 1C, and S1F**). For functional analyses, we introduced these SUZ12 mutants into K27M DMG cells expressing a doxycycline-inducible shRNA designed to knockdown endogenous *SUZ12* transcripts without affecting transgene expression.

**Figure 1.**
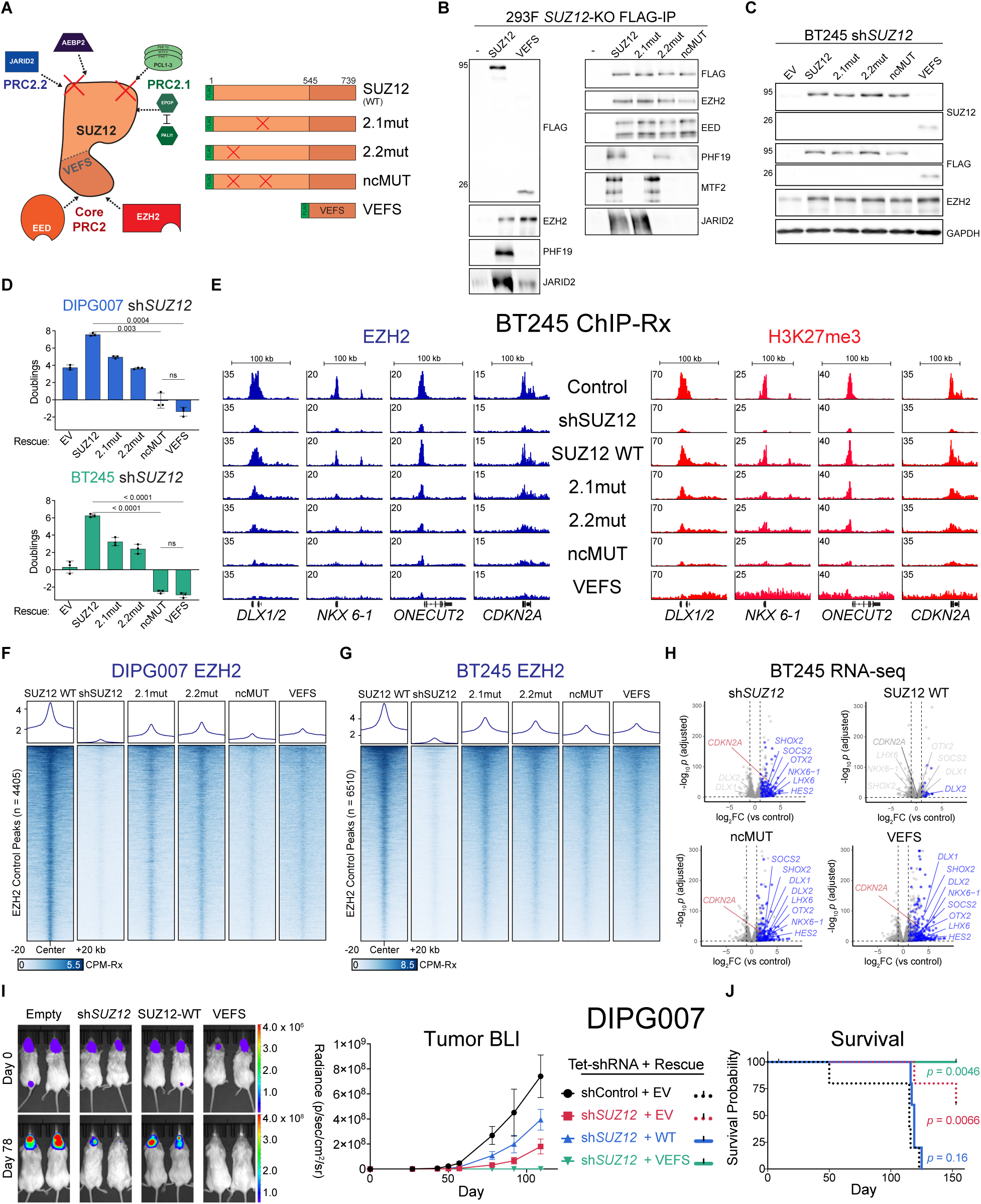
Accessory subunits collectively support K27M glioma cell proliferation through targeted PRC2 activity. **(A)** Schematic representation of SUZ12 transgenes engineered to specifically disrupt binding to PRC2 accessory subunits. All rescue constructs utilized in these experiments contained an N-terminal FLAG tag as indicated. Transgenes include: full-length wild-type SUZ12 (WT); mutants disrupting interactions with either PRC2.1 or PRC2.2 accessory subunits (2.1mut and 2.2mut, respectively); a combined non-core mutant disrupting both PRC2.1 and PRC2.2 interactions (ncMUT); and a VEFS domain-only truncation (VEFS), lacking interaction capability with all accessory subunits. **(B)** Validation of SUZ12 mutant rescue constructs by co-immunoprecipitation (co-IP) followed by immunoblot analysis. IPs were carried out in 293F SUZ12-KO cells expressing the indicated FLAG-tagged SUZ12 transgene construct. “SUZ12” indicates SUZ12 wild-type transgene rescue; “-“ indicates FLAG-negative control. Integer values to the left indicate approximate molecular weight in kDa. Antibodies used for each blot are indicated on the right. **(C)** Immunoblot analysis of whole-cell lysates from BT245 diffuse midline glioma (DMG) cells expressing a doxycycline-inducible short hairpin RNA targeting SUZ12 (Tet-shSUZ12) and constitutively expressing the indicated SUZ12 rescue transgene. Lysates were collected after 5 days of doxycycline induction. The antibody against SUZ12 detects epitopes within the VEFS domain. Molecular weights (in kDa) are indicated on the left. Cropped images for full-length SUZ12 and VEFS are from the same immunoblots and were exposed identically for accurate comparison. Antibodies used for each blot are indicated on the right. **(D)** Assessment of cell proliferation in BT245 DMG cells with SUZ12 knockdown and rescue constructs. Population doubling was quantified by cell counting after 9 days in culture. Statistical significance was determined using two-tailed t-tests with Welch’s correction; “ns” denotes not significant (p > 0.05). **(E)** Representative genome browser tracks demonstrating localization of EZH2 (upper tracks) and associated catalytic activity indicated by H3K27me3 occupancy (lower tracks) at residual genomic sites in BT245 cells under various experimental conditions. The conditions shown include: “Control,” cells expressing a non-specific short hairpin control (shControl) and an empty vector (EV); “shSUZ12,” SUZ12-depleted cells with EV; and indicated SUZ12 rescue constructs expressed in shSUZ12 cells. This labeling convention is consistently used throughout subsequent figures unless explicitly indicated otherwise. **(F-G)** Heatmaps and corresponding metaplots depicting Rx-normalized EZH2 ChIP-seq signal enrichment at peak regions identified in DIPG007 (**F**) and BT245 (**G**) control cells for each indicated experimental condition. Metaplots display the average signal intensity across these regions in reference-normalized counts per million units (CPM-Rx). **(H)** Volcano plots illustrating global transcriptome changes in BT245 cells with SUZ12 knockdown and indicated rescue constructs. Differential gene expression is normalized relative to control cells. Dashed lines indicate thresholds for statistical significance (p = 0.05) and fold change (log2 fold-change ±1). Genes highlighted in blue represent significantly upregulated transcripts proximal to PRC2 ChIP-seq peaks (detailed further in **Figure S1F**). Selected genes of biological relevance are annotated. **(I)** In vivo tumor growth assessed by bioluminescence imaging (left) of mice harboring orthotopically transplanted DIPG007 xenografts expressing indicated SUZ12 constructs, measured at defined time points post-transplantation. Quantified bioluminescence signals across all experimental conditions and time points are summarized on the right. **(J)** Kaplan-Meier survival analysis of mice depicted in panel **(J)**. Each group consists of 5 animals except the VEFS group, which includes 4 animals following exclusion of one mouse that succumbed shortly after surgery for reasons unrelated to tumor growth.

Lentiviral transduction was performed at a low multiplicity of infection (MOI) to achieve uniform, physiologically relevant transgene expression, thereby avoiding potential overexpression artifacts (**Figures 1C, S1F, and S1J**). The control condition in these knockdown/rescue experiments utilized two constructs: a non-targeting shRNA and an empty rescue vector (EV). To assess the functional importance of PRC2 subcomplex recruitment in DMG cells, we measured cell proliferation across these conditions. SUZ12 knockdown alone (EV rescue) significantly impaired cell proliferation with concurrent p16 derepression, whereas rescue with a full-length wild-type SUZ12 transgene restored proliferation and re-established p16 silencing (**Figures 1D, S1F and S1J**). Individual disruption of PRC2.1 or PRC2.2 resulted in intermediate proliferative defects, while simultaneous disruption via either the combined full-length ncMUT or the VEFS mutants severely impaired proliferation across multiple K27M DMG cell lines (**Figure 1D**). These results suggest partial redundancy and/or cooperation between PRC2.1 and PRC2.2 but strongly support a collective requirement for non-core subunits to facilitate precise PRC2 localization and functional silencing in DMG cells.

To investigate the chromatin-based mechanisms underlying the observed proliferative defects, we performed quantitative chromatin immunoprecipitation sequencing (ChIP-seq) with spike-in normalization for H3K27me3 and the PRC2 core subunit EZH2. At prominent PRC2-bound loci, including critical tumor suppressor genes such as *CDKN2A* and genes involved in neurodevelopmental pathways, both the combined full-length ncMUT and the minimal VEFS truncation constructs caused substantial reductions or complete loss of EZH2 occupancy and H3K27me3 deposition (**Figure 1E**). Conversely, loss of either PRC2.1 or PRC2.2 individually produced only intermediate reductions in both EZH2 and H3K27me3 enrichment (**Figure 1E**). More broadly, these trends held across all PRC2 recruitment sites and each K27M DMG cell line tested (**Figures 1F and 1G**). Notably, the intermediate reductions observed in 2.1mut- or 2.2mut-expressing cells did not display distinct locus-specific patterns, further supporting functional redundancy or crosstalk between PRC2.1 and PRC2.2 subcomplexes in this context. Importantly, these results strongly support the conclusion that target-site H3K27me3 represents the functionally relevant product of PRC2 catalysis in K27M DMG.

### Aberrant PRC2 recruitment rewires transcriptional networks for tumor maintenance

To elucidate how compromised PRC2 recruitment influences gene expression in K27M DMG cells, we integrated ChIP-seq for H3K27me3 and EZH2 with transcriptome-wide mRNA sequencing (RNA-seq) in the same SUZ12-mutant rescue cell lines. This integrative analysis enabled us to directly correlate changes in PRC2 chromatin occupancy with corresponding alterations in mRNA expression levels.

Relative to control cells, disruption of PRC2 localization, whether by *SUZ12* knockdown alone or by rescue with either targeting-deficient ncMUT or VEFS mutant, resulted in substantial transcriptional derepression (**Figure 1H**). Differentially upregulated genes showed significant overlap across these conditions, particularly at loci proximal to PRC2 recruitment sites identified by ChIP-seq (**Figures 1H, S1G, and S1K).**

Notably, these commonly derepressed PRC2 target genes included tumor suppressors such as *CDKN2A*, as well as genes involved in neurodevelopment, including many transcription factors with established roles in regulating neural lineage commitment and differentiation^54^ (**Figures 1H and S1M**). Gene ontology (GO) enrichment analysis of the shared gene set upregulated by impaired PRC2 recruitment revealed striking similarities with GO term categories enriched following Tazemetostat treatment across DMG models (**Figures S1I, S1L, and S1N**). These findings strongly suggest that residual, SUZ12-directed H3K27me3 maintains silencing at specific PRC2-bound loci encoding products critical for DMG pathology. Furthermore, the consistency of transcriptional responses across cell lines underscored the robustness and generalizability of our experimental models.

To determine whether the proliferative defects observed *in vitro* corresponded to impaired tumor growth *in vivo*, we employed an orthotopic mouse xenograft model using the same transgenic rescue strategy as before (**Figure 1D**). Specifically, DMG cell lines were intracranially implanted immediately after constitutive expression of SUZ12 rescue constructs, followed by induction of endogenous *SUZ12* knockdown via doxycycline-supplemented drinking water. Tumor growth was monitored longitudinally by serial bioluminescent imaging (BLI) during survival analyses (**Figure 1J**). Consistent with our cell culture findings, knockdown of endogenous *SUZ12* significantly mitigated tumor progression and improved survival compared to controls (**Figures 1I, 1J**). Remarkably, expression of the VEFS construct further reduced tumor burden and conferred a 100% survival rate. Conversely, expression of full-length wild-type SUZ12 restored tumor growth and eliminated the survival benefit associated with SUZ12 knockdown, confirming that tumor suppression was directly attributable to PRC2 ablation. Together, these results demonstrate that precise chromatin localization of PRC2, guided by the combined efforts of PRC2.1 and PRC2.2 subcomplexes, is essential for K27M DMG cell proliferation and tumorigenicity.

### The SUZ12-VEFS domain unleashes widespread PRC2 hyperactivity independent of chromatin-targeting subunits

While ncMUT and VEFS mutants successfully excluded non-core subunits from PRC2 complex incorporation (**Figure 1B**) and genomic localization (**Figures 1F-1G**) as intended, they yielded notably distinct effects on global H3K27 methylation. All full-length SUZ12 mutants, including ncMUT, restored global H3K27me3 to levels comparable to wild-type SUZ12 or parental cells, as anticipated. Unexpectedly, K27M DMG cells rescued with the VEFS truncation displayed markedly elevated bulk H3K27me3 (**Figures 2A and S1F**). These observations contrast with prior studies in mouse and human embryonic stem cells (mESCs/hESCs), in which similar genetic manipulations aimed at disrupting PRC2.1 and PRC2.2 subcomplexes did not significantly alter global H3K27 methylation levels, including experiments utilizing VEFS rescue in SUZ12-KO mESCs^39,48,52^. To determine whether VEFS-induced increases in H3K27me3 were unique to K27M DMG cells, we generated SUZ12-knockout mouse embryonic fibroblasts (MEFs), which express only wild-type histone H3, and performed rescue experiments using the same SUZ12 transgenes. Consistent with the DMG cell data, VEFS-rescued MEFs similarly exhibited increased global H3K27me3, whereas full-length SUZ12 mutants restored methylation to wild-type levels (**Figure 2B**). These patterns, initially identified by immunoblotting, were independently confirmed using quantitative mass spectrometry-based histone modification profiling (**Figure 2C**). To further assess the broader applicability of these findings, we examined whether VEFS-containing PRC2 could overcome inhibition by EZHIP, an endogenous H3 K27M mutation. Specifically, we introduced a dTAG degron^55^ into the EZHIP locus of two EZHIP-expressing cell lines, followed by SUZ12 knockout and subsequent rescue with full-length wild-type SUZ12, ncMUT, or VEFS transgenes. Mirroring findings from DMG cells and MEFs, VEFS-rescued cells exhibited significantly elevated H3K27me3 compared to ncMUT- or wild-type SUZ12-rescued counterparts under both inhibitory (EZHIP-expressing) and permissive (EZHIP-degraded) conditions (**Figures S2A, S2B**).

**Figure 2.**
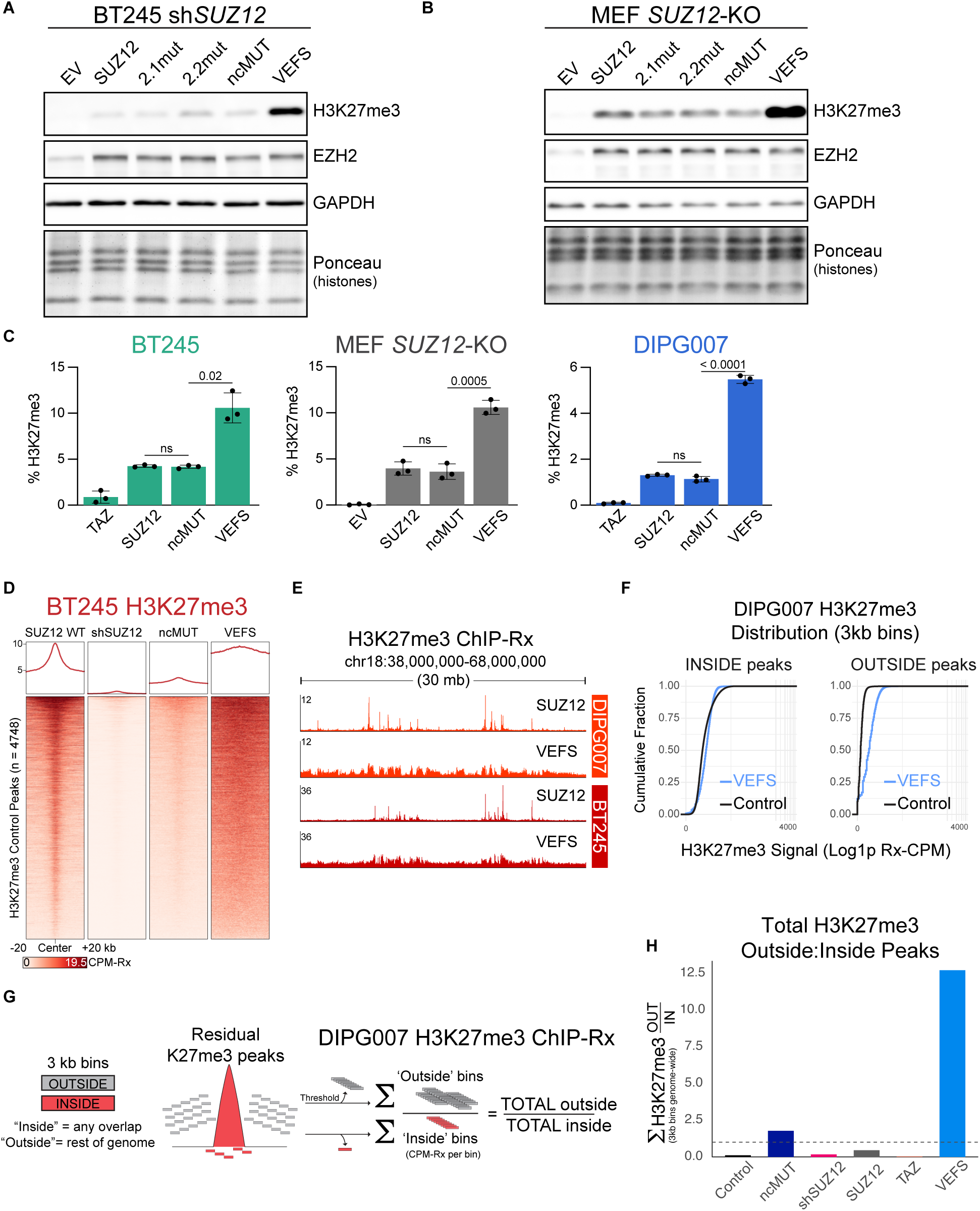
VEFS-expressing cells globally increase H3K27me3 independent of accessory subunits. **(A)** Immunoblot analysis of whole-cell lysates from BT245 rescue samples. Antibodies used for each blot are indicated on the right; ponceau staining is included as an additional loading control. **(B)** Immunoblot analysis of whole-cell lysates from SUZ12 knockout (KO) MEFs rescued with the indicated SUZ12 transgenes. Antibodies used for each blot are indicated on the right; ponceau staining is included as an additional loading control. **(C)** Quantitative analysis of H3K27me3 levels by histone PTM mass spectrometry in SUZ12 knockdown (DIPG007/BT245) or knockout (MEF) cell lines rescued with the indicated SUZ12 transgenes. Negative controls were included for comparison: for DIPG007/BT245, control cells (shControl + EV) were exposed to 5 µM Tazemetostat for 9 days, while MEF SUZ12-KO transduced with an empty vector were used as a negative control. Percent H3K27me3 values represent aggregate measurements of all H3 isoforms. Histones were extracted from the same chromatin pellets used for ChIP-seq **(2E, 2F, and S2B)**. Displayed p values derived from two-tailed t-tests with Welch’s correction (n = 3 biological replicates per condition). **(D)** Spike-in normalized ChIP-seq (ChIP-Rx) H3K27me3 signal in BT245 cells expressing the indicated SUZ12 rescue constructs. Heatmaps and associated metaplots are centered on peaks called in control cells. Metaplots display the average signal intensity across these regions in reference-normalized counts per million units (CPM-Rx). **(E)** Genome browser views of a 30mb region on chr18, highlighting diffuse elevations in H3K27me3 consistent across DMG cell lines. **(F)** Empirical cumulative distribution function (ECDF) plots showing the genome-wide distribution of H3K27me3 ChIP-seq signal across 3 kb bins in control versus VEFS-rescue samples. Bins are stratified by whether they overlap control-defined H3K27me3 peaks (“inside”) or fall outside peak regions (“outside”), illustrating the extent to which VEFS-driven H3K27me3 is redistributed into previously unmarked genomic regions. **(G)** Schematic representation for genome-wide analyses of H3K27me3 ChIP-Rx data. Bins were generated and quantified for data in **Figures 2F, 2H, S2F, and S2G**. Peak-overlapping bins are depicted as red blocks while all other bins falling outside the limits of residual peaks are depicted in gray. The basic approach to quantifying total levels of signal in these regions relates to **Figures 2H** and **S2G** specifically. **(H)** Bar plot showing the relative distribution of total H3K27me3 ChIP-seq signal outside versus inside residual PRC2 peaks (as defined in control DIPG007 cells). Approach to this analysis is described in the above legend for **Figure 2G**. The dotted line corresponds to a value of 1.0; proportions above this represent cell states with more total H3K27me3 outside of peak regions than within.

The SUZ12 N-terminus forms a regulatory lobe structurally distinct from the catalytic lobe of PRC2 and is widely recognized for its role as a scaffold facilitating interactions with auxiliary subunits that modulate PRC2 function. However, our observation that the VEFS mutant globally elevates H3K27me3 levels compared to the ncMUT mutant suggests an additional regulatory function intrinsic to the N-terminus itself, independently of auxiliary subunit recruitment. This represents a mechanism that, to our knowledge, has not been described. To investigate how VEFS enhances PRC2 activity, we extended our H3K27me3 ChIP-seq analyses to map the genomic distribution of this modification in VEFS-rescued K27M DMG cells. Across all residual peaks identified in control cells, average H3K27me3 signals in VEFS-rescued DMG cells were markedly increased compared to cells rescued with ncMUT or other full-length SUZ12 mutants (**Figures 2D, S2C, and S2D**), despite both VEFS and ncMUT mutants displaying similarly impaired genomic localization (**Figures 1F and 1G**). Notably, this observation contrasts with previous reports in VEFS-rescued SUZ12-KO mESCs, in which PRC2 occupancy and H3K27me3 deposition at recruitment sites were diminished. To confirm that these results were not confounded by incomplete knockdown of endogenous SUZ12, we performed parallel ChIP-seq experiments assessing H3K27me3 and EZH2 distribution in SUZ12-KO MEF rescues. Similar to our findings in K27M DMG cells, VEFS rescue in SUZ12-KO MEFs also resulted in substantial elevations of H3K27me3 signal at control loci compared to ncMUT, despite both mutant forms exhibiting markedly compromised PRC2 occupancy at canonical sites (**Figure S2E**).

Although VEFS increased the average H3K27me3 signal at control peaks, the nature of this elevation was notably flattened compared to wild-type (**Figures 2D, S2C-S2E**). At a genome-wide level, VEFS cells showed broad, contiguous accumulation of H3K27me3 that spanned megabase-sized regions, contrasting sharply with the punctate enrichments seen in wild-type SUZ12 DMG cells (**Figure 2E**). To characterize these diffuse H3K27me3 patterns quantitatively, we segmented the genome into uniform segments (3 kb bins), measured the average H3K27me3 ChIP-Rx signal within each bin, and classified these bins based as either overlapping with control-defined peaks (“inside peaks”) or located outside these peaks (“outside peaks”).

Cumulative distribution analysis of H3K27me3 (**Figure 2F**) showed that within target sites, VEFS modestly enriched low- and mid-intensity bins, consistent with the uniform elevation seen in averaged metaplots (**Figures 2D, S2C, and S2D**). In contrast, the top ∼10% of peak bins in control cells had higher H3K27me3 ChIP-Rx values than any VEFS bins, consistent with our earlier observation that VEFS fails to produce high levels of H3K27me3 at strong loci such as *CDKN2A* (**Figures 2F and 1E**). Moreover, differences in H3K27me3 enrichment were far more pronounced for bins outside PRC2 peaks (**Figure 2F**). In control cells, virtually all inter-peak bins lacked detectable methylation, consistent with K27M-mediated confinement of H3K27me3 in DMG. In VEFS cells, however, over half of all inter-peak bins acquired low-to-moderate H3K27me3. Since inter-peak regions span ∼95% of the genome, these data suggest VEFS-mediated hypermethylation observed by bulk analyses reflects widespread, low-level methylation across large genomic domains rather than from enhanced methylation at canonical PRC2 target sites.

We next investigated whether these broadly distributed but weakly methylated genomic regions generated by VEFS could effectively engage downstream reader complexes. To approximate the pool of H3K27me3 available to these reader proteins, we calculated the total H3K27me3 signal by summing across all qualifying 3 kb bins within each bin category and compared their relative contributions in each experimental condition (**Figure 2G**). We employed a conservative threshold to exclude binds with minimal signal, ensuring noise-driven data were omitted. Remarkably, in VEFS-expressing cells, most of the total H3K27me3 signal was found in genomic regions outside canonical PRC2 target loci, significantly surpassing the aggregate intensity within canonical peaks (**Figure 2H**). In contrast, nearly all detectable H3K27me3 in K27M cells rescued with full-length SUZ12 remained tightly confined to canonical PRC2-bound sites. Importantly, these findings were consistent across a broad range of signal filtering thresholds, underscoring their biological robustness and potential significance (**Figure S2G**).

Together, these data reveal an unexpected intrinsic function of the SUZ12 N-terminal domain in restraining global PRC2 catalytic activity, likely independent of auxiliary subunit interactions. Our quantitative genome-wide analyses demonstrated that VEFS-expressing cells, lacking this regulatory constraint, generated widespread, diffuse domains of H3K27me3 deposition spanning extensive genomic regions. Although the per-base intensity of H3K27me3 within these broad domains was relatively low, their massive genomic coverage significantly altered the overall cellular distribution of H3K27me3. Such extensive, diffuse methylation could plausibly divert downstream reader complexes away from their intended high-affinity target loci, potentially impacting cellular transcriptional regulation.

### Diffuse H3K27me3 deposition induced by VEFS displaces canonical PRC1 from its genomic targets

Traditionally, Polycomb-mediated gene repression involves recruitment of canonical PRC1 (cPRC1) complexes to H3K27me3-enriched chromatin, primarily mediated through chromodomain-containing CBX proteins^10^. This recruitment facilitates chromatin compaction and transcriptional silencing^56^. To characterize the impact of VEFS-induced H3K27me3 redistribution on cPRC1 localization, we performed spike-in ChIP-seq for canonical PRC1 subunits CBX4 and BMI1 (also known as PCGF4), both of which have been implicated in DMG ^57–61^. To disentangle the effects of PRC2 localization, targeted activity, and diffuse methylation on cPRC1 recruitment, we compared the following four DMG conditions: (1) control cells with intact PRC2, (2) Tazemetostat-treated cells with intact but catalytically-inhibited PRC2, (3) VEFS-rescued cells with impaired targeting but globally elevated H3K27me3 levels, and (4) *SUZ12* knockdown cells deficient in both PRC2 recruitment and catalysis.

cPRC1 binding sites were defined as the combined peaks of CBX4 and BMI1 in control cells (**Figure S3A**), which extensively overlapped with residual PRC2-bound loci (**Figure S3E**). As expected, both Tazemetostat treatment and *SUZ12* knockdown reduced H3K27me3 and cPRC1 binding; however, this effect varied considerably across individual loci. For example, *KLF9* showed complete loss of cPRC1 occupancy, *LHX1* retained partial occupancy, and *DLX1/2* remained largely occupied despite K27me3 ablation (**Figures 3A and 3B**). Notably, VEFS-expressing cells displayed near-complete loss of cPRC1 occupancy across these same sites (**Figures 3A and 3B**). These locus-specific trends held true across cPRC1 peaks. Treatment with Tazemetostat or SUZ12 knockdown resulted in modest reductions in cPRC1 localization, whereas VEFS expression led to markedly more severe loss of cPRC1 localization, despite generating higher levels of H3K27me3 at these same sites (**Figures 3C-3F**). This uncoupling of cPRC1 occupancy from H3K27me3 deposition was particularly evident at *HOX* loci, where substantial levels of cPRC1 enrichment persisted in Tazemetostat-treated and *SUZ12*-knockdown cells despite extensive depletion of H3K27me3 and PRC2 localization (**Figures 3E and S3E**). In contrast, VEFS-expressing cells exhibited pronounced disruption of these prominent Polycomb domains, resulting in near-complete loss of cPRC1 subunit occupancy across each HOX cluster. These observations were consistent across DMG cell models, PRC1 subunits, ChIP quantification methods, and biological replicates (**Figures 3C, 3D, S3B, S3C, and S3D**).

**Figure 3.**
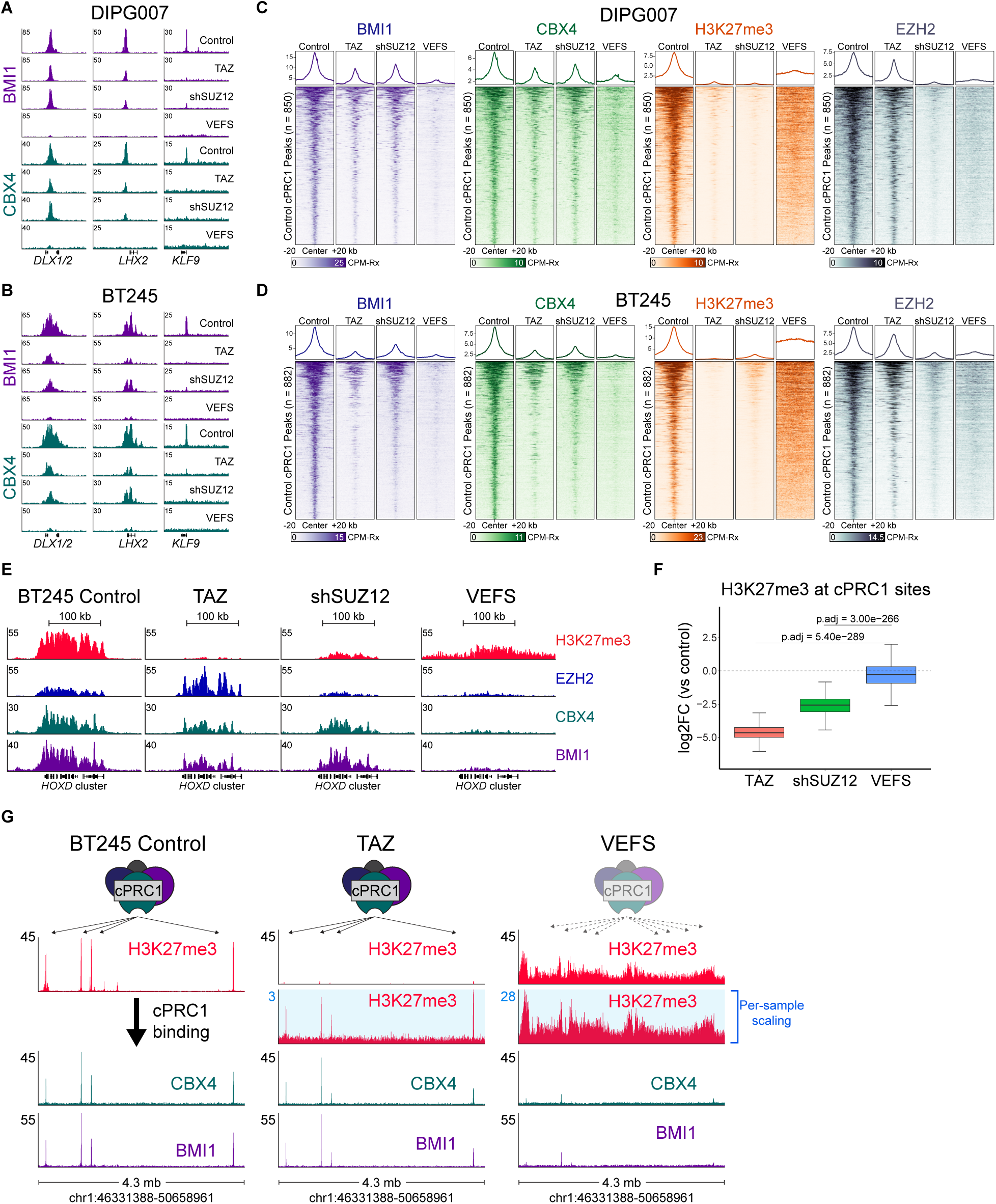
Diffuse H3K27me3 Elevation Displaces Canonical PRC1 More Effectively Than PRC2 Loss. **(A-B)** Genome browser tracks displaying ChIP-seq signal for canonical PRC1 subunits BMI1 and CBX4 at loci encoding regulators of neurodevelopment. DIPG007 (**A**) and BT245 (**B**) samples with the indicated conditions are shown on the right of each track. **(C-D)** Heatmaps and corresponding metaplots showing Rx-normalized ChIP-seq signal for BMI1, CBX4, H3K27me3, and EZH2 centered on canonical PRC1 (cPRC1) sites. These sites are defined as merged overlaps of BMI1 and CBX4 peaks called in control cells (see **Figure S3A**). Results are shown separately for DIPG007 (**C**) and BT245 (**D**). **(E)** Genome browser views of ChIP-Rx signal for H3K27me3, EZH2, CBX4, and BMI1 in BT245 cells at the *HOXD* gene cluster, representative of densely-populated polycomb sites. See **Figure S3E** for similar visualization of *HOXC*. *HOXA* and *HOXB* clusters (not shown) exhibit similar patterns and levels of enrichment across all conditions **(F)** Boxplot quantification of log₂ fold-change in H3K27me3 ChIP-seq signal intensity in BT245 cells at cPRC1 peaks defined in **(S3A)**. Note that the underlying data represents the metaplots in **Figure 3D** but is shown here to emphasize decoupling average enrichment values irrespective of distribution across peaks. p-values are pairwise comparisons derived from Wilcoxon rank-sum tests with FDR correction. **(G)** Genome browser visualization of ChIP-seq signal for canonical PRC1 subunits (CBX4, BMI1) and H3K27me3 across a representative 4.3 Mb region on chromosome 1. The bottom two cPRC1 and top H3K27me3 tracks are scaled uniformly across conditions (Control, Tazemetostat, and VEFS) for cross-sample comparison. The blue-highlighted H3K27me3 tracks are auto-scaled to emphasize signal dynamics in low-enrichment contexts, particularly the diminished H3K27me3 signal in Tazemetostat-treated cells. PRC1 cartoons depict relationships between H3K27me3 distributions and cPRC1 localization. The dashed arrows and faded transparency in the case of VEFS depicts non-productive localization as a result of diffuse H3K27me3 distributions.

To better understand the differences between VEFS expression and loss of PRC2 activity in directing PRC1 localization, we examined a 4 Mb region containing multiple prominent PRC2 target loci (**Figure 3G**). Control cells exhibited tight correspondence between narrow domains of residual H3K27me3 and sharply defined cPRC1 occupancy. Similarly, Tazemetostat-treated cells retained robust cPRC1 enrichment at these loci despite nearly undetectable total H3K27me3 signals by ChIP-Rx normalization. However, autoscaled genome browser views revealed that the minimal residual methylation in Tazemetostat-treated cells remained selectively concentrated at canonical peaks, maintaining a favorable local signal-to-noise ratio sufficient to recruit PRC1. In contrast, VEFS expression resulted in broad distribution of H3K27me3 throughout surrounding genomic regions, diluting the relative prominence of high-affinity PRC1-binding sites.

Collectively, our findings demonstrate that VEFS expression induces a global redistribution of H3K27me3 that differs markedly in both magnitude and spatial distribution from scenarios involving PRC2 depletion. The observation that VEFS expression more effectively displaces canonical PRC1 compared to either catalytic inhibition or complex depletion challenges the conventional model in which cPRC1 localizes to chromatin predominantly via instructive H3K27me3 patterns. Instead, our data support a model wherein diffuse methylation functions as a molecular decoy, actively titrating cPRC1 complexes away from their canonical binding sites and effectively destabilizing Polycomb domains. Although accumulating evidence has indicated analogous mechanisms of chromatin targeting in other cellular contexts^62^, the functional relevance of nonspecific H3K27me3 patterns for determining cPRC1 occupancy remains largely unexplored^63–66^. Given the uniquely confined landscape of H3K27me3 in DMG cells, disruptions in this spatial equilibrium may represent a distinct therapeutic vulnerability in K27M-driven gliomas.

### VEFS-driven chromatin hypermethylation dominantly impairs DMG cell proliferation

The ability of VEFS to broadly redistribute H3K27me3 and displace canonical PRC1 in rescue experiments suggested it might exert dominant-negative effects on cell fitness in the context of intact endogenous SUZ12. Supporting this possibility, preliminary experiments revealed that VEFS expression alone (without doxycycline-induced knockdown) impaired cell proliferation and induced p16 (**Figures S1J and S4A**). To confirm these observations and explore the underlying molecular processes, we ectopically expressed VEFS or control constructs without knockdown of endogenous *SUZ12* mRNA, again utilizing lentiviral transductions at low MOI. To confirm that VEFS-induced effects were specifically due to PRC2 interactions rather than off-target toxicity, we generated VEFS mutants designed to disrupt interactions with core PRC2 subunits. Initially, we tested the N618Y mutation, previously shown in *Drosophila* to abate PRC2 subunit interactions, but this alone was insufficient in mammalian cells (**Figures S4B and S4C**). Therefore, we introduced an additional mutation, S662R, predicted to sterically impair the SUZ12–EZH2 interaction interface within the VEFS domain (**Figure S4B**). This double mutant (NY.SR) successfully abolished PRC2 complex formation, confirmed via co-immunoprecipitation (**Figure S4C**).

Subsequent proliferation assays demonstrated that wild-type VEFS robustly inhibited growth across all tested DMG cell lines, whereas the PRC2 interaction-deficient NY.SR mutant exhibited no growth-inhibitory effect (**Figures 4A–4D**). Importantly, K27M DMG cell clones lacking p16 expression were resistant to VEFS-mediated growth inhibition, consistent with their previously demonstrated resistance to PRC2 inhibitor Tazemetostat (**Figures S1D and S1E**). In contrast, glioblastoma cells expressing wild-type histone H3 (GBM002) were unaffected by VEFS expression, paralleling their resistance to PRC2 inhibitors (**Figure S1B**). Together, these results indicate that VEFS and Tazemetostat share a common mechanism of inhibiting DMG cell proliferation through dysregulation of PRC2-dependent gene silencing, despite their distinct molecular activities.

**Figure 4.**
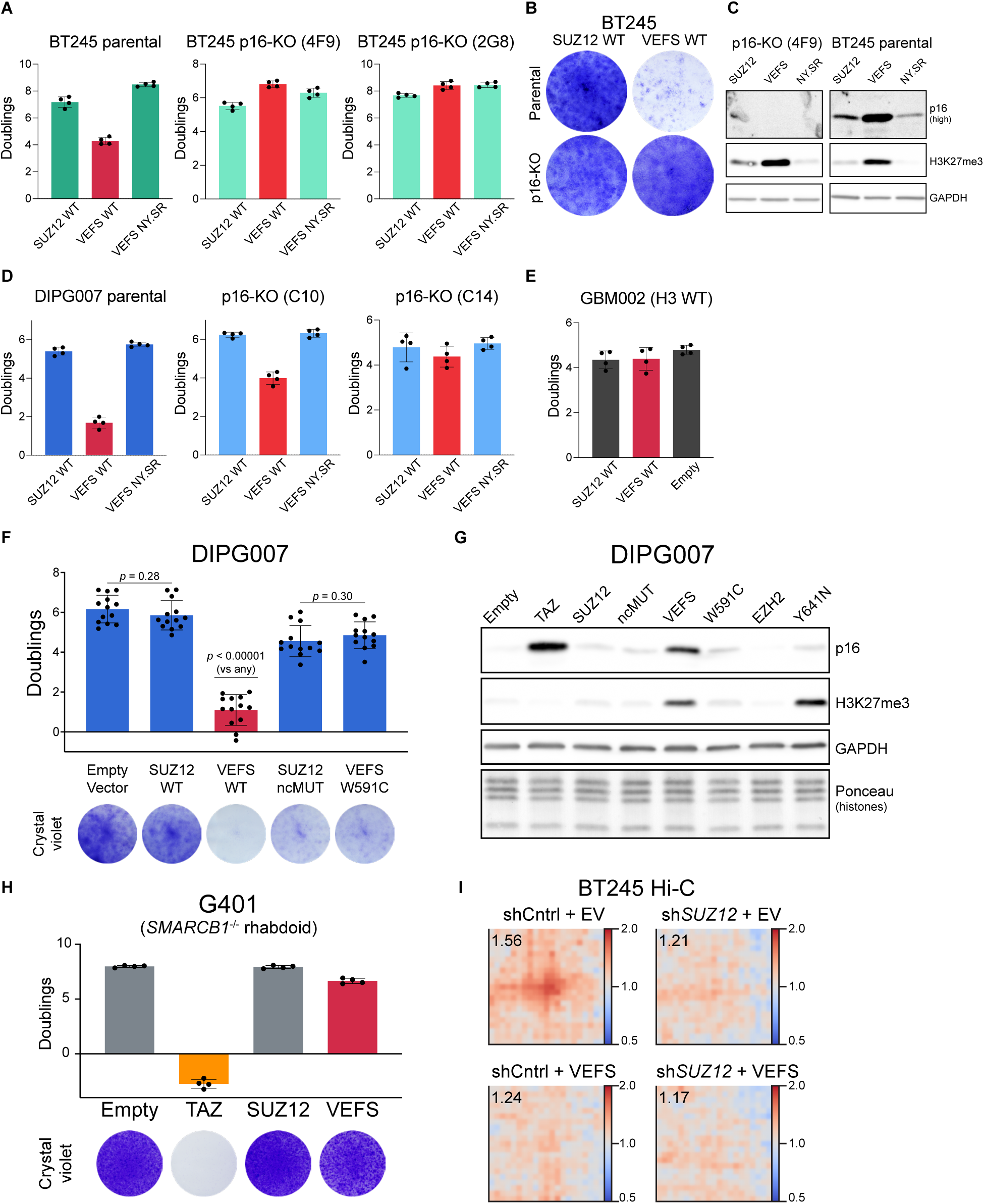
VEFS Domain Expression Alone Dominantly Impairs DIPG Cell Proliferation by Inducing Diffuse Gains in Cellular H3K27me3. Note that all data represented here, with the exception of select data in 4K (as noted in-figure), are derived from cells ectopically expressing the indicated constructs in the context of intact endogenous SUZ12 (no knockdown with rescue as in prior figures). **(A)** Proliferation assays in BT245 DMG cells ectopically expressing the indicated SUZ12 constructs. These include: full-length wild-type SUZ12 (SUZ12 WT), VEFS-only truncation (VEFS WT), and the VEFS truncation harboring the N618Y/S662R double mutation (NY.SR VEFS), which disrupts core PRC2 complex formation (see Supplemental Figure for validation). Proliferation was also assessed in two clonal BT245 lines (4F9, 4G8) with homozygous deletion of *p16^6INK4A^* (see **Figure S1**) Data represent mean population doublings in 9-day experiments from four biological replicates across two independent experiments. **(B)** Representative images of crystal violet-stained monolayers corresponding to samples in panel **(A)**. **(C)** Immunoblots of whole-cell lysates from BT245 parental and p16 knockout clones transduced with the indicated SUZ12 constructs. High-exposure anti-p16 blots are included to highlight successful knockout, H3K27me3 blots highlight consistent hypermethylation for VEFS-expressing samples regardless of p16 status, and GAPDH blots are included as loading controls. **(D)** Proliferation assays in parental and p16-KO DIPG007 clones (C10, C14) expressing the indicated SUZ12 constructs. Experimental conditions parallel those in **(A)**. Each condition was assessed with four biological replicates in two independent experiments. **(E)** Proliferation assay in an H3 wild-type pediatric glioblastoma cell expressing the indicated SUZ12 constructs. Experimental conditions parallel those in **(A)** and **(D)** and again represent four biological replicates across two independent experiments. **(F)** Proliferation assays in DIPG007 cells expressing the following SUZ12 constructs: full-length wild-type (SUZ12-WT), VEFS truncation (VEFS-WT), full-length non-core mutant (ncMUT), and catalytically inactive VEFS-W591C mutant. The top panel shows population doublings over a 9-day period (n = 13 biological replicates from 4 independent experiments). The bottom panel shows representative crystal violet-stained monolayers from matched cultures. Displayed p-values from two-tailed t-tests with Welch’s correction. ns, not significant (p > 0.05). **(G)** Immunoblots of whole-cell lysates from DIPG007 cells expressing the same SUZ12 transgenes defined in **(F)** or EZH2 transgenes (“EZH2” refers to wild-type, “Y641N” is an EZH2 gain-of-function mutant). Note “Empty” and “TAZ” samples were transduced with an empty vector control with the latter receiving Tazemetostat treatment (5 µM). Lysates were harvested 9 days after transduction or drug exposure. **(H)** Proliferation assays in G401 rhabdoid tumor cells (which lack the SWI/SNF core subunit *SMARCB1*) expressing the indicated SUZ12 constructs. A Tazemetostat-treated group transduced with an empty vector is included for comparison. Top panel shows population doublings assessed on day 9 (n = 4 from 2 independent experiments); bottom panel shows representative crystal violet-stained monolayers. **(I)** Hi-C pile-up plots showing pairwise chromatin interaction frequency at regions enriched for both CBX4 and H3K27me3 (as defined by ChIP peaks in BT245 controls). Each plot reflects the average interaction frequency (observed/expected) for locus pairs in a given condition. Top left: untreated control; top right: SUZ12 knockdown alone; bottom right: SUZ12 knockdown with VEFS rescue; bottom left: VEFS expression alone (in the presence of endogenous SUZ12). Enrichment in the center of each plot represents punctate, long-range interactions anchored at Polycomb-bound regions, which are partially retained or reorganized depending on the degree and nature of H3K27me3 spreading.

To elucidate the molecular mechanisms underlying VEFS-induced cytotoxicity, we assessed proliferation in DMG cells expressing various SUZ12 constructs. We initially hypothesized that VEFS might bind PRC2 core subunits (EZH2/EED), interfering with endogenous PRC2 assembly. Supporting this idea, ncMUT and VEFS conferred similarly lethal consequences for K27M DMG cells (**Figure 1D**). Additionally, ncMUT expression in cells with intact SUZ12 modestly reduced proliferation compared to controls (**Figure 4F**), suggesting competition with endogenous SUZ12 for incorporation into PRC2. Alternatively, the smaller VEFS truncation may integrate more efficiently into PRC2 complexes than full-length ncMUT, thus preferentially disrupting endogenous PRC2 complexes.

Alternatively, this difference may reflect a unique gain-of-function activity seen in VEFS but not ncMUT, specifically its ability to generate widespread, diffuse H3K27me3 deposition and actively displace PRC1 from its normal genomic targets (**Figures 2, S2, and 3**). To disentangle out-competition due to protein size from VEFS catalytic gains, we used a VEFS mutant, W591C, known to disrupt EZH2 catalytic function. Critically, expression of the catalytic mutant VEFS truncation caused only a mild growth disadvantage, nearly identical in magnitude to that seen in cells expressing ncMUT (**Figure 4F**). These results strongly suggest that the dominant-negative phenotypes associated with VEFS expression primarily result from catalytic hyperactivation.

To further pinpoint the mechanism underlying VEFS-induced toxicity, we hypothesized that its detrimental effects might stem specifically from altered spatial distribution (diffuse accumulation) rather than increased overall abundance of H3K27me3. To test this hypothesis, we sought to elevate global H3K27me3 levels without disrupting the focal, SUZ12-directed recruitment of PRC2. To this end, we expressed the hyperactive EZH2 mutant Y641N, which increases catalytic activity but is not predicted to affect SUZ12-directed recruitment. Indeed, expression of Y641N successfully elevated H3K27me3 to levels similar to VEFS but did not impact proliferation or induce p16 expression (**Figures 4I and S4E**). Consistent with this observation, prior ChIP-seq analyses confirmed that Y641N retained focal PRC2 localization at canonical targets sites (**Figure S4D**)^67^. Together, these results strongly suggest that VEFS-mediated toxicity specifically arises from diffuse rather than focal H3K27me3 deposition.

Given the distinct, CpG island-confined pattern of H3K27me3 in K27M DMG, we hypothesized that sensitivity to diffuse methylation may vary in other PRC2-dependent cancers. To evaluate this possibility, we expressed VEFS and appropriate controls in *SMARCB1*-deficient rhabdoid tumor cells (G401), a PRC2-dependent cell line known for broader enhancer-associated H3K27me3 distribution^68^. In line with our hypothesis, G401 cells were largely resistant to VEFS-driven proliferation defects, despite showing pronounced sensitivity to H3K27me3 loss upon treatment with Tazemetostat (**Figures 4H, S4F**). These findings reinforce our model that spatial distribution, rather than overall abundance of H3K27me3, dictates cell-type-specific sensitivity to aberrant PRC2 activity.

Recent insights assessing 3D genomic organization in K27M DMG revealed a critical role for cPRC1 in organizing higher-order chromatin structures to maintain oncogenic gene expression^61,69^. To assess the impact of VEFS hypermethylation on 3D chromatin organization, we performed chromatin conformation capture (Hi-C) in VEFS-expressing DMG cells. To measure cPRC1-dependent chromatin looping, we quantified pairwise contact frequencies between distant genomic regions centered on cPRC1 ChIP-seq sites (**Figure 4K**). These analyses demonstrated that VEFS expression alone significantly altered cPRC1 chromatin looping, reducing central observed-over-expected (O/E) contact values from 1.56 in controls to 1.24, closely matching the disruption observed with SUZ12 knockdown alone (O/E = 1.21) (**Figure 4I**). The combination of VEFS expression and SUZ12 depletion produced the most pronounced chromatin disruption (O/E = 1.17), emphasizing the potent effects of VEFS on chromatin structures and reinforcing its dominant effects in DMG cells.

Together, these findings reveal that VEFS-driven hypermethylation dominantly impairs DMG cell proliferation through widespread, diffuse H3K27me3 deposition and disruption of cPRC1-dependent chromatin organization. However, the precise molecular determinants within SUZ12 that normally prevent such global hyperactivity remained unclear.

### The SUZ12 N-terminus constrains PRC2 catalysis through nucleic acid interactions

To define structural regions in the SUZ12 N-terminus responsible for restricting PRC2 activity in cells, we expressed a series of SUZ12 truncation constructs in MEF SUZ12 knockout cells and analyzed global H3K27me3 levels (**Figure 5A**). Informed by high-resolution PRC2 structures, we found that inclusion of the WDB2 domain immediately N-terminal to VEFS in the “NEC” truncation construct (named by the first three amino acids, starting with N492) resulted in elevated H3K27me3 levels comparable to those seen with VEFS expression (**Figure 5B**). In contrast, inclusion of N-terminal regions beyond this domain decreased H3K27me3 to full-length levels, suggesting that the region spanning R427 (start of the “RIF” truncation) and N492 negatively regulates PRC2 catalysis in cells.

**Figure 5.**
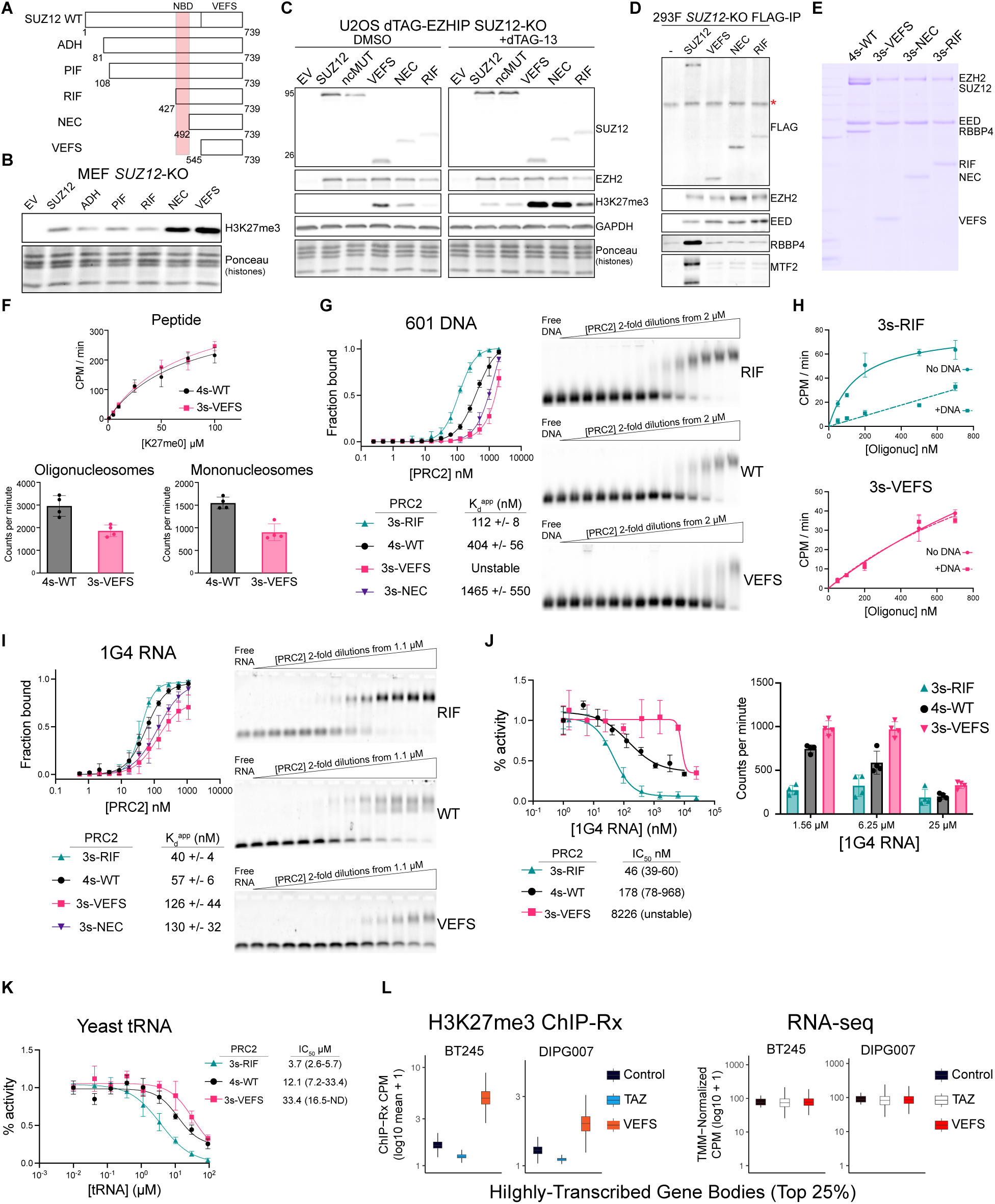
A Discrete Region in the N-Terminus of SUZ12 Restrains Global PRC2 Activity by Limiting Indiscriminate Nucleic Acid Interactions. **(A)** Schematic representation of a panel of SUZ12 truncation constructs designed to map the minimal region required to prevent aberrant H3K27me3 accumulation. Each construct is named by the first three amino acids encoded at its N-terminus, with numbering corresponding to their position in the full-length canonical SUZ12 isoform. **(B)** Immunoblot analysis of whole-cell lysates from SUZ12-knockout MEFs expressing the indicated SUZ12 truncation constructs. Antibodies used for each blot are indicated on the right; ponceau staining is included as a loading control. **(C)** Immunoblot analysis of whole-cell lysates from SUZ12-knockout U2OS cells expressing dTAG-degradable EZHIP and the indicated SUZ12 truncation constructs. Where indicated, dTAG-13 treatment (500 nM for 6 days) was used to degrade EZHIP protein. Antibodies used for each blot are indicated on the right; ponceau staining is included as an additional loading control. **(D)** Co-immunoprecipitation assays performed in SUZ12-knockout cells to test whether truncation constructs can interact with PRC2 core and accessory subunits. FLAG-tagged SUZ12 truncations were immunoprecipitated, and eluates were probed by immunoblotting. A red asterisk indicates a non-specific band that was also present in the no-FLAG control. Antibodies used for each blot are indicated on the right - EZH2, EED, and RBBP4 are core subunits and MTF2 is an accessory subunit. **(E)** Coomassie-stained SDS-PAGE gel showing purified PRC2 complexes used for biochemical assays in this figure. “4s” (four-subunit complex) includes EZH2, EED, RBBP4, and wild-type SUZ12. “3s” complexes consist of EZH2, EED, and the indicated truncated forms of SUZ12. **(F)** Baseline histone methyltransferase activity assays comparing wild-type and truncated PRC2 complexes. Assays were performed using H3 tail peptides (tested across a titration series), mononucleosomes, or native oligonucleosomes (each at 200 nM). Reactions were conducted at 20 nM PRC2 in 40 mM KCl for 45 minutes. Methylation activity was measured by ^3^H-SAM incorporation and expressed as counts per minute (CPM). **(G)** PRC2 binding to nonspecific free DNA (Widom 601 sequence) assessed by EMSA. Left: quantification of DNA-protein complex formation using a Cy5-labeled probe, averaged across three independent experiments. Right: representative EMSA gels. **(H)** Inhibition of PRC2 HMT activity by free Widom 601 DNA. Activity was measured in reactions containing oligonucleosomes across a range of concentrations, with or without 1 µM free 601 DNA. Reactions were carried out with 20 nM PRC2 at 40 mM KCl for 30 minutes. CPM, (scintillation) counts per minute corrected for background. **(I)** EMSA measuring binding of PRC2 complexes to a synthetic RNA oligonucleotide containing a single G-quadruplex-forming motif (1G4). Left: quantification of gel shifts from three independent experiments. Right: representative agarose gels. **(J)** HMT inhibition assays using the same G-quadruplex RNA (1G4) tested in panel (I). Reactions were performed with 40 nM PRC2 and 400 nM native oligonucleosomes in 100 mM KCl for 30 minutes. Left: activity normalized to RNA-free reactions, shown as percent activity (as determined by scintillation) for IC₅₀ calculation. Right: background-corrected CPM values from the same reactions, allowing direct comparison across PRC2 complexes. **(K)** HMT inhibition assays using yeast tRNA to assess general RNA-mediated inhibition. Percent activity and IC₅₀ values were determined by scintillation as in panel **(J)**, under identical reaction conditions. **(L)** Boxplots summarizing ChIP-seq and RNA-seq quantifications across highly transcribed genes in DMG cells. The genes included in these analyses represent the top 25% most highly expressed transcripts (as determined by RNA-seq counts in control cells for each cell line). Left: average H3K27me3 ChIP-seq signal spanning gene bodies; right: TMM-normalized RNA-seq expression values shown as counts per million (CPM).

To determine if these observations apply broadly across different cellular contexts, including those with and without expression of the K27M-like inhibitor EZHIP, we introduced the same SUZ12 truncations into a SUZ12-knockout dTAG-EZHIP cell line. Consistent with our findings in MEFs, expression of either VEFS or NEC truncations resulted in marked elevations in global H3K27me3, whereas expression of RIF or full-length SUZ12 did not alter methylation levels significantly (**Figure 5C**).

The region between RIF and NEC encompasses a relatively uncharacterized zinc finger domain; its role in binding nucleic acids and how these events affect PRC2 function have not, to our knowledge, been rigorously examined. Although previous structural analyses identified this region as beta sheet-rich and situated immediately C-terminal to a known RRM-like domain^23,70^, the functional significance of nucleic acid binding within this region, including its implications for PRC2 regulation, has not yet been extensively examined. Notably, recent cryo-electron microscopy studies revealed direct points of contact between residues in this zinc finger-containing region and linker DNA^71^, suggesting a functional role in DNA engagement. Based on these insights and our findings described below, we refer to this region as the nucleic acid binding domain (NBD) (**Figure 5A**).

Previous structural work also identified contact points between this Zn finger region and accessory subunits, such as JARID2, in reconstituted systems^42^. However, we hypothesized RIF would fail to interact with full-length accessory subunits in cells based on our earlier manipulations lying far N-terminal to R427 (**Figures 1A and 1B**). Additionally, it was unclear from available structural data whether RBBP4/7 proteins would associate with RIF or NEC truncations. Co-immunoprecipitation experiments carried out in *SUZ12*-KO cells showed that all three truncations of interest (VEFS, NEC, and RIF) assembled minimal catalytic core complexes with EZH2 and EED, but failed to associate with accessory subunits or RBBP4 (**Figure 5D**). Thus, these constructs enabled us to specifically investigate the intrinsic function of the SUZ12 zinc finger-containing region independently of its role as a scaffold for accessory subunits. To this end, we purified three-subunit assemblies (3s-VEFS, 3s-NEC, and 3s-RIF) and a four-subunit control complex (4s-WT), containing RBBP4 and full-length wild-type SUZ12, to represent native core PRC2 (**Figure 5E**).

To directly assess the functional consequences of SUZ12 truncations, we purified PRC2 complexes from HEK293 and Sf9 cells and performed histone methyltransferase (HMT) assays on various substrates (**Figures 5E, 5F, S5B, S5C**). VEFS complexes (3s-VEFS) displayed comparable catalytic activity on isolated H3K27 peptides comparable to full-length core PRC2 (4s-WT). However, 3s-VEFS showed consistently reduced activity on chromatin substrates, such as mononucleosomes and native oligonucleosomes (**Figures 5F, S5A-D**). These findings indicate that VEFS-driven hypermethylation observed in cells does not reflect enhanced intrinsic catalytic potential but rather indicates a distinct gain-of-function phenotype arising specifically in the cellular context.

To test whether nucleic acids regulate PRC2 activity through interactions with the SUZ12 NBD, we conducted binding assays (electrophoretic mobility shift assays, EMSAs) and HMT activity assays using oligonucleosome substrates in the presence or absence of exogenous nucleic acids. Initially, using Widom 601 DNA as a nonspecific dsDNA control, we found NBD-containing complexes (4s-WT, 3s-RIF) bound DNA with nanomolar affinity, whereas complexes lacking this domain (3s-VEFS and 3s-NEC) exhibited substantially reduced affinity (**Figure 5G**). In functional assays, free DNA potently inhibited nucleosome-directed catalytic activity of 3s-RIF complexes, whereas 3s-VEFS complexes were unaffected (**Figure 5H**).

Similar experiments conducted with a short RNA oligonucleotide capable of forming a G-quadruplex, previously reported to bind and inhibit PRC2^70^, demonstrated strong binding to NBD-containing complexes (3s-RIF, 4s-WT), but significantly weaker binding to 3s-VEFS and 3s-NEC complexes (**Figure 5I**). Functional assays confirmed potent RNA-mediated inhibition of RIF and full-length PRC2, whereas VEFS-containing complexes remained largely active even at high RNA concentrations (**Figure 5J**). To further confirm the generality of these findings, we repeated these assays using G-quadruplex forming DNA oligonucleotide^72^ which similarly displayed reduced affinity for NBD-lacking complexes (**Figure S5E**). Additionally, using unfolded yeast tRNA as a generalized RNA inhibitor, we observed approximately a 10-fold increase in the IC_50_ for 3s-VEFS compared to NBD-containing 3s-RIF complexes (**Figure 5K**). Together, these results indicate the SUZ12 NBD region binds diverse nucleic acid species with high affinity and that these interactions significantly inhibit histone methyltransferase activity *in vitro*.

In cellular contexts, transcription is known to preclude H3K27me3 through several proposed mechanisms, including RNA-mediated inhibition of PRC2 catalysis^10,73^. Thus, we hypothesized that VEFS might be aberrantly active in transcribed regions due to its lack of NBD-based regulation. To this end, we identified highly-expressed genes (top 25% expression by RNA-seq) in control DMG cells and quantified average H3K27me3 signal across these gene bodies. Consistent with our predictions, VEFS expression led to impressive H3K27me3 accumulations across highly-transcribed gene bodies, presumably within RNA-rich nuclear environments typically hospitable to PRC2 catalysis. Despite considerable increases in H3K27me3 in these regions, we observed no apparent gene-silencing effects locally (**Figure 5L**), supporting our earlier conclusions (**Figure 3**) that the blankets of H3K27me3 deposited by PRC2-VEFS primarily impact gene silencing through diversion of PRC1 away from target sites rather than establishing new repression of nearby genes. Together, these findings suggest that VEFS-expressing cells lacking the NBD-RNA interactions responsible for inhibiting PRC2 activity in our reconstituted biochemical models correlated with unchecked PRC2 catalysis in nuclear regions rich in these same components.

## Discussion

Our findings provide mechanistic insight into the regulation of PRC2 enzymatic activity, identifying the SUZ12 N-terminal region as a modulatory hub that constrains PRC2 through transient interactions with nucleic acids mediated in part by the nucleic acid binding domain (NBD). We find that the dynamic interplay between productive nucleosome engagement and non-productive nucleic acid binding establishes a kinetic equilibrium essential for precise chromatin targeting. Specifically, truncation of the SUZ12 N-terminal regulatory domain, including the NBD, disrupts this equilibrium (**Figure 6A**), resulting in hyperactive, indiscriminate deposition of H3K27me3, displacement of canonical PRC1 complexes, perturbation of three-dimensional chromatin architecture, and impairment of tumor growth in H3 K27M-driven DMGs (**Figure 6B**). These results suggest that Polycomb specificity is governed, at least in part, by a kinetic buffering process, enhancing our understanding of PRC2 regulation and pointing to potential therapeutic vulnerabilities in PRC2-dependent malignancies.

**Figure 6.**
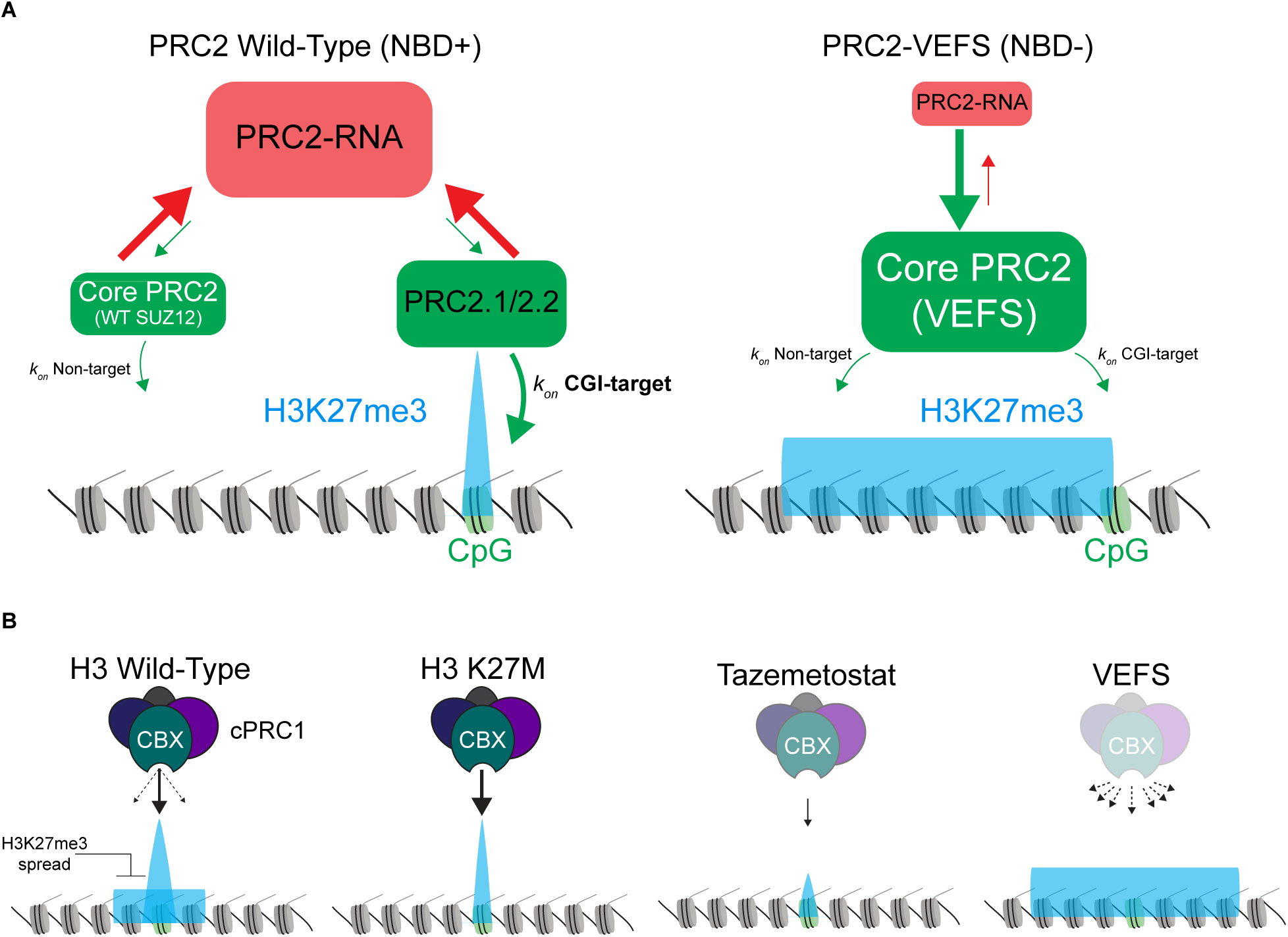
Model for SUZ12 NBD-RNA Interactions in Establishing Equilibria Necessary for Targeted Polycomb Silencing in Cells. **(A)** Proposed kinetic equilibrium model illustrating how the SUZ12 N-terminal nucleic acid-binding domain (NBD) dynamically regulates PRC2 activity through transient interactions with RNA, thereby constraining its catalytic function to appropriate genomic targets and preventing widespread methylation at inappropriate non-target sites. Arrow thicknesses represent approximate relative transition rates between RNA-bound and RNA-free states, and the relative sizes of PRC2 cartoons reflect the steady-state abundance of each complex within cells. Global H3K27me3 distribution patterns are depicted in blue, illustrating targeted versus widespread methylation under these distinct regulatory conditions. **Left panel** (PRC2 Wild-Type, NBD+): Under physiological conditions, the presence of the SUZ12 NBD allows both core PRC2 (comprising EZH2, EED, and SUZ12) and the larger PRC2.1/2.2 complexes to extensively and transiently associate with cellular RNA (depicted in red). These RNA-PRC2 interactions are predominantly inhibitory, generating catalytically inactive species and limiting PRC2 methyltransferase activity. Due to the high abundance and promiscuous nature of RNA binding, the equilibrium strongly favors inactive RNA-bound PRC2, minimizing off-target histone methylation. Only complexes harboring accessory subunits with a high affinity for CpG island (CGI) chromatin targets (green nucleosomes) significantly overcome these RNA-mediated inhibitory interactions, shifting the equilibrium towards an RNA-free, catalytically active state (green), thus achieving precise, targeted deposition of H3K27me3 (blue peak). In this scenario, widespread promiscuous RNA interactions act as a kinetic buffer to ensure PRC2 specificity by restricting methylation to designated genomic sites. **Right panel** (PRC2-VEFS, NBD−): In contrast, when the critical N-terminal RNA-binding region of SUZ12 is deleted or absent, as exemplified by the VEFS-containing truncated PRC2 complex, RNA-mediated inhibition is greatly diminished. Without this regulatory interaction, the equilibrium shifts markedly towards a state favoring RNA-free, catalytically competent PRC2 complexes. Consequently, this leads to widespread, indiscriminate chromatin binding and methylation activity at both canonical CGI-targets and non-target genomic sites, represented by a diffuse and extensive pattern of H3K27me3 enrichment across multiple nucleosomes. **(B)** Schematic modeling the downstream consequences for canonical PRC1 (cPRC1) readers when H3K27me3 patterns are altered. H3K27me3 distributions are again represented in blue and CGI target sites are represented as a green nucleosome. cPRC1 assemblies localize to polycomb domains via a CBX paralogue binding high local concentrations of PRC2-deposited H3K27me3 (depicted as solid arrows, binding strength reflected by arrow thickness). In cells, cPRC1 binding strength is refined by low-affinity, transient interactions with poorly concentrated H3K27me3 (depicted as dashed arrows) outside of polycomb sites. In typical H3 wild-type cells, this balance is established by PRC2 spreading from sites of initial recruitment. In DMG cells, K27M inhibits PRC2 spread, resulting in a restricted pattern of H3K27me3 that confines PRC1 to CGI targets to maintain oncogenic gene silencing. This dependency can be exploited by reducing target-site H3K27me3 (via PRC2-ablative strategies such as Tazemetostat) or by increasing the total pool of H3K27me3 elsewhere to titrate PRC1 away, as we observed in cells expressing VEFS. In our cell models, this latter approach resulted in more potent reductions in PRC1 binding (**Figure 3**), providing rationale to investigate therapies that simulate VEFS-driven hypermethylation.

### Kinetic Equilibrium Governs PRC2 Substrate Specificity

Previous biochemical analyses have established that PRC2 exhibits markedly lower intrinsic affinity for its nucleosomal substrates, typically in the micromolar range, compared to its substantially higher nanomolar affinity for various nucleic acids, including RNA and DNA ^74–79^. This pronounced difference in affinities has presented a paradox: how PRC2 achieves precise and selective genomic targeting despite its binding preference favoring abundant nonspecific nucleic acid substrates over specific nucleosomal substrates^16^.

We propose that this paradox is resolved by kinetic buffering, a regulatory mechanism in which enzyme specificity and activity are governed by transient interactions with abundant nonspecific substrates. This principle has been broadly observed across multiple biochemical systems, including transcription factors^80–83^, DNA and RNA polymerases^84,85^, and protein kinases^86,87^. In the context of PRC2, kinetic buffering occurs through transient, nonproductive interactions with abundant nucleic acids, effectively limiting the complex from engaging nonspecifically with substrate nucleosomes. Our biochemical and genomic analyses of the SUZ12 N-terminal domains strongly support this model (**Figure 6A**), demonstrating that these domains mediate non-productive nucleic acid interactions, thereby restricting PRC2 enzymatic activity by limiting productive nucleosome engagement. Recent evidence further supports the concept of kinetic buffer, indicating that PRC2 binding to abundant, nonspecific RNAs effectively reduces inappropriate enzymatic activity and enhancing genomic targeting specificity ^88,89^.

This kinetic model is further substantiated by the biochemical and genomic effects observed upon isolated expression of the VEFS domain, which produced hyperactive PRC2 complexes characterized by reduced nucleic acid affinity and elevated enzymatic activity independent of canonical chromatin-recruitment mechanisms. Notably, kinetic buffering mediated by transient nucleic acid interactions complements and expands existing frameworks of PRC2 regulation, including the well-characterized allosteric activation pathway involving the EED subunit^17^. While the allosteric mechanism enhances catalytic activity at specific genomic loci through positive feedback loops driven by pre-existing H3K27me3 marks, kinetic buffering via nonspecific nucleic acid interactions imposes a broader constraint on PRC2 activity. Consequently, these two regulatory mechanisms function concurrently, precisely calibrating PRC2 activity in response to local chromatin states and nucleoplasmic environments.

Building upon previous studies identifying PRC2 interactions with DNA and diverse RNA species, we highlight the functional significance of structured RNAs, particularly those forming G-quadruplex motifs. These structured RNAs competitively inhibit PRC2 interactions with substrate nucleosomes, thereby constraining catalytic activity and refining genomic targeting precision. Additionally, specific long noncoding RNAs, such as Xist and HOTAIR, actively recruit PRC2 to defined genomic loci, thereby balancing locus-specific gene repression against generalized catalytic activity. Together, these diverse nucleic acid interactions maintain PRC2 in a dynamic kinetic equilibrium, precisely modulating substrate specificity, genomic localization, and overall epigenomic stability.

Collectively, our findings establish that PRC2 specificity and activity are governed not by static binding alone but rather by a finely tuned kinetic equilibrium that balances productive nucleosome interactions against non-productive nucleic acid engagement. This equilibrium is critically modulated by structural domains within SUZ12, providing novel mechanistic insights into the precise regulation of chromatin-modifying activities essential for proper gene expression control during development and implicated in disease pathology.

### SUZ12 N-terminal Domains Regulate PRC2 Activity

Our detailed biochemical analyses demonstrate that the SUZ12 N-terminal domains act as critical regulatory hubs within PRC2 (**Figure 6A**), actively constraining its enzymatic output through transient interactions with nucleic acids. Specifically, we have identified that removal of these N-terminal regulatory elements, encompassing a previously uncharacterized zinc finger domain now designated as the nucleic acid binding domain (NBD), significantly alters PRC2 functionality. Complexes lacking the N-terminal regulatory region exhibit reduced affinity for nucleic acids, leading to unrestrained catalytic activity, and reduced sensitivity to H3 K27M and EZHIP inhibitors. These data indicate that the SUZ12 N-terminal domains function not merely as a structural scaffold, but as an essential molecular gatekeeper, precisely regulating PRC2 enzymatic activity and genomic targeting specificity.

Previous studies have predominantly identified EZH2 as the principal RNA-binding subunit within PRC2, supported by biochemical and structural evidence demonstrating direct interactions between structured RNAs and residues distributed throughout EZH2 domains^75,79,90,91^. Complementing this body of work, additional reports have also implicated SUZ12 in RNA binding^74,77^. Our findings expand upon these studies by specifically dissecting the regulatory role of SUZ12 N-terminal region independently of EZH2-mediated interactions. Our biochemical experiments reveal that the NBD within SUZ12 binds diverse nucleic acids, including structured RNAs and DNA, with high affinity, effectively restricting access of PRC2 complexes to nucleosomal substrates under normal cellular conditions. This interaction establishes a kinetic buffering mechanism, constraining PRC2 catalytic activity and enforcing precise chromatin targeting.

### Intergenic Chromatin as a Molecular Sink Regulating Polycomb Repression

Our study provides insights into the regulatory capacity of intergenic chromatin, highlighting its role as a molecular sink that dynamically modulates the genomic availability and targeting of PRC2. Historically viewed as inert, intergenic regions have emerged as active modulators of chromatin states^92–94^. Our findings demonstrate that extensive stretches of intergenic chromatin impede PRC2 through numerous nonspecific binding events, establishing a kinetic equilibrium that limits excessive enzymatic activity and ensures the targeted, focal deposition of H3K27me3. This spatial buffering thereby prevents aberrant methylation within gene-rich regulatory regions, preserving the fidelity of Polycomb-mediated repression.

The buffering role of intergenic chromatin is supported by our experiments employing the ncMUT and VEFS mutants. Expression of the VEFS construct results in PRC2 hyperactivation, causing widespread, diffuse H3K27me3 deposition across intergenic regions, consistent with disruption of the equilibrium depicted in **Figure 6A**. This broad, non-focal methylation redistributes cPRC1 occupancy away from targeted developmental loci, as cPRC1 requires concentrated, focal H3K27me3 domains for stable binding and repression^56^. Consequently, despite elevated global H3K27me3 levels, cPRC1 occupancy is significantly diminished at critical Polycomb target genes, including *CDKN2A* and key neurodevelopmental regulators, resulting in paradoxical transcriptional derepression. These findings underscore that focal H3K27me3 enrichment is essential for effective Polycomb repression through PRC1-driven chromatin compaction and transcriptional inhibition^10^.

Our high-resolution chromatin conformation capture (Hi-C) experiments further underscore the necessity of focalized H3K27me3 deposition. Diffuse methylation patterns resulting from VEFS expression compromise normal chromatin interactions and disrupt genomic compartmentalization, emphasizing that precise chromatin modifications are critical for maintaining the three-dimensional architecture required for transcriptional repression. These observations underscore an intricate interplay between histone modifications and higher-order chromatin organization in the regulation of gene expression.

In contrast to VEFS, expression of ncMUT disrupts recruitment of specific accessory subunits that are required for precise chromatin localization but does not substantially the catalytic activity of PRC2. In cells expressing ncMUT, reductions in H3K27me3 deposition and subsequent cPRC1 occupancy occur selectively at canonical Polycomb target loci, without significant global redistribution. Thus, although both VEFS and ncMUT mutants impair cPRC1 occupancy, their underlying mechanisms differ markedly: ncMUT induces localized deregulation, whereas VEFS drives widespread H3K27me3 and titration of cPRC1 into intergenic regions.

Recognizing intergenic chromatin as an active component of chromatin regulation expands our understanding of genomic dynamics and transcriptional precision. Disruption of intergenic buffering capacity, whether through elevated enzymatic activity, impaired recruitment of accessory subunits, or altered chromatin interactions, can lead to aberrant gene expression and developmental dysfunction. This concept is reflected in other chromatin regulatory contexts; for example, loss of BAP1 inducing global enrichment of H2AK119ub1, which redistributes PRC1 and PRC2 away from canonical regulatory targets^64–66^. Similarly, mutations such as H3 K36M decrease H3K36me2/3 levels, resulting in ectopic deposition of PRC2 and H3K27me3 and widespread dispersion of PRC1 binding^63^. An analogous phenomenon occurs in yeast, where loss of DOT1 and H3K79me3 leads to titration of the silencing SIR2/3/4 complex, disrupting gene repression due to limited factor availability at specific loci ^62^.

Collectively, these examples illustrate a conserved regulatory theme wherein aberrant redistribution of histone modifications disperses effector complexes, thereby impairing their ability to concentrate effectively at critical regulatory loci. By characterizing intergenic chromatin as a regulatory buffer, our findings uncover previously unappreciated aspects of chromatin biology that carry potential therapeutic implications, particularly in conditions involving Polycomb misregulation.

### Clinical Implications of PRC2 Regulatory Mechanisms in Cancer

Our findings provide critical insights into the clinical implications of PRC2 regulation in cancer, particularly in DMGs characterized by the H3 K27M mutation or aberrant EZHIP expression. Previous literature has suggested a significant role for PRC2 activity in DMG proliferation, although discrepancies regarding its exact functional contribution persist. Our results demonstrate that residual PRC2 catalytic activity remains essential for tumor cell proliferation in H3 K27M-mutant DMGs, supporting the hypothesis that complete PRC2 inhibition may represent an efficacious therapeutic approach for this malignancy.

Although substantial evidence implicates PRC2 dysregulation in various malignancies^95,96^, clinical outcomes from therapies targeting core PRC2 components, notably EZH2, have exhibited significant variability, showing efficacy in specific lymphoma subtypes but yielding limited success in solid tumors^97–99^. Our findings suggest that exploiting the nuanced regulatory dynamics of PRC2 complexes, particularly the kinetic buffering mechanisms mediated by transient interactions with nucleic acids, could enhance therapeutic precision. For instance, our finding that widespread and aberrant H3K27me3 deposition, induced by VEFS domain expression, disrupts cPRC1 occupancy and perturbs tumor-specific chromatin architecture identifies a potential therapeutic vulnerability. Specifically, inducing hyperactive but mislocalized PRC2 activity, as modeled by VEFS expression (**Figure 6B**), could disrupt pathological chromatin configurations, resulting in transcriptional derepression of tumor suppressor genes and significantly inhibiting tumor cell growth.

Moreover, our findings highlight an essential role for accessory subunit-guided recruitment of PRC2 in K27M DMG. While previously reported redundancy or synergy among subcomplexes theoretically complicates direct clinical translation, our findings provide important clarification over conclusions drawn from another recent study^61^. That study found that loss of individual accessory non-core subunits did not significantly affect clonal fitness in a pooled CRISPR dropout screen, whereas gRNAs targeting core PRC2 members were enriched, leading to the assertion that non-core subunits lack relevance in DMG pathology. Although not inconsistent with these prior observations, our results distinctly emphasize the functional importance of PRC2.1/2.2-instructed recruitment mechanisms in this disease. Clarifying these regulatory details is essential for precisely defining the molecular underpinnings of DMG and may have clinical relevance if future therapeutic approaches effectively target entire classes of accessory subcomplexes.

In summary, our work clarifies essential regulatory mechanisms governing PRC2 activity, emphasizing kinetic buffering as a critical determinant of enzymatic specificity and genomic targeting fidelity. These insights present substantial opportunities for the development of innovative therapeutic strategies. An integrated understanding of PRC2 subunit dynamics, nucleic acid interactions, and chromatin regulatory functions will be essential for designing targeted interventions. Such strategies, specifically directed at disrupting oncogenic chromatin states, represent promising avenues for therapeutic advancement.

## Method Details

### Cell Culture and Lentivirus Production

Glioma cells were cultured as monolayers on tissue culture plates coated with laminin at a concentration of 1 μg/cm². Cells were maintained in DMEM/F12 basal media supplemented with B-27 minus vitamin A (Gibco), N-2 supplement (Gibco), heparin (2 μg/mL), epidermal growth factor (EGF, 20 ng/mL), fibroblast growth factor (FGF, 10 ng/mL), L-alanyl-L-glutamine (2 mM), and antibiotic-antimycotic solution (1X).

Non-glioma cell lines were cultured in DMEM with high glucose, supplemented with 10% fetal bovine serum (FBS), L-alanyl-L-glutamine (2 mM), and penicillin-streptomycin (1X). Cells were maintained as adherent monolayers. All cell lines were routinely tested and confirmed to be free of mycoplasma contamination.

Lentiviral particles were produced using HEK293T cells transfected with VSV-G and psPAX2 packaging plasmids along with either a pCDH or pLKO-Tet transfer plasmids. Viral supernatants were collected at 24 and 48 hours post-transfection and concentrated by adding 1/3 volume of 4X homemade lenti concentrator (20% PEG-6000, 20% PEG-8000, 760 mM NaCl, 1.6X PBS) and incubating at 4C overnight. Concentrated virus was pelleted by centrifugation at 1500×g for 45 minutes and resuspended in DMEM/F12 at 1/100th the original supernatant volume.

### Lentivirus Titer Quantification

Lentiviral titers were determined separately for each cell line. Cells were transduced with a 3-fold dilution series of concentrated virus. At 24 hours post-transduction, monolayers were split into four wells: two with antibiotic (at a minimal concentration needed for 100% cell death) and two without antibiotic. In parallel, untransduced control cells were plated under identical conditions. Cell viability was monitored, and once untransduced cells in the antibiotic-treated wells were fully non-viable, the corresponding transduced wells were harvested and counted. Only conditions yielding 5–30% viability relative to non-antibiotic controls were used to calculate viral titer, expressed as transducing units per milliliter (TU/mL).

### Generation of Transgenic SUZ12 Cell Lines

Adherent monolayer cells were transduced with constitutive pCDH-based lentiviruses in the presence of 5 μg/mL polybrene. Virus was applied at a multiplicity of infection (MOI) of approximately 0.2 to favor single-copy genomic integration. Transduced cells were selected with 7.5 μg/mL blasticidin for 4–5 days. Prior to use in downstream functional assays, all samples were passaged once post-selection at a standardized sub-confluent seeding density. For rescue experiments, all cell lines (including controls) contained Tet-shRNA constructs previously selected for with 3 µg/mL puromycin but were not exposed to doxycycline prior to start of the experiment.

### Cell proliferation assays

Cells were seeded at a density of 7,000 cells per well in 12-well plates. At designated time points, cells were harvested using Accutase and counted using a CytoSMART automated cell counter. In replating assays, cells were reseeded at the same density (7,000 cells per well) at defined intervals, which were selected based on preliminary experiments to correspond with sub-confluent densities in control or untreated conditions. For crystal violet staining, parallel wells were seeded at identical densities in 6-well plates. At the time of cell counting, monolayers were fixed with 4% paraformaldehyde in PBS for 10 minutes at room temperature, then stained with 0.5% crystal violet for 10 minutes. Excess stain was removed, and wells were rinsed thoroughly with Milli-Q water. Plates were dried overnight and imaged using a flatbed scanner.

### Chromatin Immunoprecipitation

40-60 million cells were cross-linked in 10 mL of 1% paraformaldehyde (PFA) in PBS for 10 minutes at room temperature with end-over-end rotation. Reactions were quenched by adding glycine to a final concentration of 200 mM and rotating for an additional 5 minutes. Cells were pelleted at 500 × g for 5 minutes at 4 °C, washed twice with 10 mL PBS, flash-frozen, and stored at −80 °C.

For chromatin extraction, cross-linked cell pellets were resuspended in Lysis Buffer I (50 mM HEPES pH 7.9, 140 mM NaCl, 1 mM EDTA, 10% glycerol, 0.5% NP-40, 0.25% Triton X-100) supplemented with 1X protease inhibitor cocktail (Pierce, EDTA-free), and incubated for 10 minutes at 4 °C with rotation. Nuclei were pelleted at 1,400 × g for 5 minutes and washed twice with MNase buffer (50 mM HEPES pH 7.9, 1 mM CaCl₂, 20 mM NaCl, 0.5 mM PMSF, 1X PIC). The chromatin pellet was resuspended in 950 μL MNase buffer and digested with micrococcal nuclease (Worthington) at 37 °C for 10 minutes. MNase units were empirically optimized for each cell line and crosslinking condition to achieve primarily mononucleosomal fragments. The reaction was quenched by the addition of 10 mM EDTA, 5 mM EGTA, 80 mM NaCl, 0.1% sodium deoxycholate, and 0.5% sodium lauroyl sarcosine.

Chromatin further treated with light sonication using a Covaris system (PIP 140, duty factor 5%, cycles per burst 200; 3 cycles of 45 seconds on, 30 seconds off). Triton X-100 was added to a final concentration of 1%, and the sample was clarified by centrifugation at 21,000 x g for 10 minutes at 4C. The supernatant was dialyzed for 2 hours at 4C against RIPA buffer (10 mM Tris-HCl pH 7.6, 1 mM EDTA, 0.1% SDS, 0.1% sodium deoxycholate, 1% Triton X-100, 150 mM NaCl, 0.5 mM PMSF), using 3.5 kDa MWCO snakeskin dialysis tubing. Following dialysis, chromatin was clarified again by centrifugation at 21,000 x g for 10 minutes. Glycerol was added to a final concentration of 10% before flash-freezing. Chromatin concentrations were measured by qubit high-sensitivity dsDNA assay.

For immunoprecipitation chromatin samples were mixed with ∼1% (by weight) of spike-in chromatin in 500 µl of ChIP binding buffer (RIPA buffer with 150 mM NaCl and 0.8 µM PMSF). For histone ChIPs, drosophila (S2 cell) chromatin was used for spike-in. For PRC2/PRC1 subunit ChIPs in human glioma samples, MEF chromatin was used for spike-in. For EZH2 ChIP in MEF samples, HEK293F chromatin was used for exogenous spike-in. 0.5-2 µg of antibody were used per 5-10 µg of sample chromatin and incubated with rotation at 4C overnight.

Chromatin bound to antibody was captured by incubating with anti-mouse or anti-rabbit Dynabeads at 4C for 3 hours and then washed: 3x with RIPA + 150 mM NaCl, 2x with RIPA + 300 mM NaCl, 2x with LiCl buffer (0.25 M LiCl, 0.5% NP-40, 0.5% Na-DOC), and 1x with TE buffer + 50 mM NaCl. Each wash consisted of 10-minute incubations with end-over-end rotation at 4C. Finally, DNA was eluted in ChIP elution buffer (10 mM Tris, 1 mM EDTA, 1% SDS) and crosslinks reversed overnight by incubation at 65C with agitation. Eluted material was treated with RNase A (1 μL, 37 °C for 1 hour) and proteinase K (1 μL, 55 °C for 2 hours), followed by purification using a PCR column-based DNA purification kit (IBI), and eluted in 60 μL TE buffer. Sequencing libraries were prepared using NEBNext Ultra II DNA library kits according to the manufacturer’s instructions.

Libraries were sequenced on an Illumina NovaSeq X Plus or HiSeq 4000 1×50bp at approximately 20×10^6^ reads per sample.

### Murine tumor models

All animal studies were approved by the Institutional Animal Care and Use Committee (IACUC) at the University of Wisconsin – Madison. Female Nod-Rag-Gamma (NRG; Jax #007799) mice (NRG) were purchased at 4 to 5 weeks of age from The Jackson Laboratory (jax.org) and used for all experiments. Mouse experiments were performed with at least four animals per treatment group in each trial; aggregate number of animals (n) is indicated.

### Xenograft experiments

Intracranial implantation of DMG cells was performed as previously described^100^. Briefly, cancer cells were enzymatically dissociated to single cells (2×10^5^ cells) and suspended in 5 µl of cell culture medium. Using a Hamilton syringe under aseptic conditions in a biosafety cabinet, the cells were stereotactically injected into the Pons of anesthetized NRG mice at 1 µl/min at the following coordinates referenced from bregma: -4.84 mm antero-posterior, +1 mm medio-lateral, and -4.5 mm dorso-ventral. At specific time-points or at onset of neurological symptoms, tumor-bearing mice were euthanized, and brains excised and processed for further analyses. Doxycycline was administered to mice via drinking water at concentration of 2 mg/ml. Water consumption by mice was monitored routinely, and fresh doxycycline-containing water was prepared and changed weekly.

### Bioluminescent Imaging

For bioluminescent whole body imaging, mice were administered D-luciferin (150 mg/kg, i.p.) at 10-12 minutes prior to imaging. Mice were anesthetized using isoflurane through a nose cone and placed into the imager (IVIS Spectrum; Perkin Elmer). Care was taken to loosely place mice into nose cone to avoid background noise from nose cone onto location of tumor cell injection in head. Imaging times were optimized by manufacturers software (Living Image; Revvity Inc.) to achieve optimal signal acquisition for all groups of mice. Post-imaging, bioluminescent signal was analyzed using manufacturers software (Living Image; Revvity Inc.) by drawing region-of-interest to include entire mouse head, and analyzed data plotted as photon flux (photons/second/m^2^).

### Acid Extraction of Histones for LC-MS/MS analysis

Approximately 8 × 10⁶ DIPG cells were pelleted, washed with PBS, flash-frozen, and stored at −80 °C. Thawed pellets were lysed in hypotonic buffer (20 mM HEPES pH 7.9, 10 mM KCl, 2 mM MgCl₂, 0.1 mM EDTA, 0.5 mM EGTA, 1 mM benzamidine, 0.8 mM PMSF, 1 mM DTT) and incubated for 30 minutes at 4 °C. Nuclei were pelleted and extracted in 0.4 N H₂SO₄ for 6 hours with rotation. Debris was cleared by centrifugation, and histones were precipitated with TCA (25% final), incubated overnight on ice, and washed with cold acetone (±0.1% HCl). Pellets were air-dried, resuspended in water, clarified by centrifugation, and stored at −80C. Extracted histones were prepared for chemical derivatization and digestion as described previously^101,102^. In brief, the lysine residues from histones were derivatized with the propionylation reagent (1:2 reagent:sample ratio) containing acetonitrile and propionic anhydride (3:1), and the solution pH was adjusted to 8.0 using ammonium hydroxide. The propionylation was performed twice and the samples were dried on speed vac. The derivatized histones were then digested with trypsin at a 1:50 ratio (wt/wt) in 50 mM ammonium bicarbonate buffer at 37°C overnight. The N-termini of histone peptides were derivatized with the propionylation reagent twice and dried on speed vac. The peptides were desalted with the self-packed C18 stage tip. The purified peptides were then dried and reconstituted in 0.1% formic acid. An LC-MS/MS system consisted of a Vanquish Neo UHPLC coupled to an Orbitrap Exploris 240 (Thermo Scientific) was used for peptide analysis. Histones peptide samples were maintained at 7 °C on sample tray in LC. Separation of peptides was carried out on an Easy-Spray™ PepMap™ Neo nano-column (2 µm, C18, 75 µm X 150 mm) at room temperature with a mobile phase. The chromatography conditions consisted of a linear gradient from 2 to 32% solvent B (0.1% formic acid in 100% acetonitrile) in solvent A (0.1% formic acid in water) over 48 min and then 42 to 98% solvent B over 12 min at a flow rate of 300 nL/min. The mass spectrometer was programmed for data-independent acquisition (DIA). One acquisition cycle consisted of a full MS scan, 35 DIA MS/MS scans of 24 m/z isolation width starting from 295 m/z to reach 1100 m/z. Typically, full MS scans were acquired in the Orbitrap mass analyzer across 290–1100 m/z at a resolution of 60,000 in positive profile mode with an auto maximum injection time and an AGC target of 300%. MS/MS data from HCD fragmentation was collected in the the Orbitrap. These scans typically used an NCE of 30, an AGC target of 1000%, and a maximum injection time of 60 ms. Histone MS data were analyzed with EpiProfile^103^.

### Purification of native oligonucleosomes from human cells

Native oligonucleosomes were purified from SUZ12-KO 293F cells. Nuclei were prepared by resuspending cell pellets in hypotonic lysis buffer A (20 mM HEPES pH 7.9, 10 mM KCl, 2 mM MgCl_2_, 0.1 mM EDTA, and 0.5 mM EGTA supplemented with 1 mM benzamidine, 0.8 mM PMSF, 1 mM DTT, 0.5 mM spermine, and 0.15 mM spermidine) followed by douncing. Nuclei were pelleted and washed once with buffer A + 0.25% triton X-100 to clean up cytoplasmic material. Soluble nuclear proteins were extracted by resuspending nuclei in buffer C (0.42 M NaCl, 25% glycerol, 0.2 mM EDTA supplemented with 0.5 mM spermine, 0.15 mM spermidine, 1X PIC, 0.4 mM PMSF, and 1 mM DTT), followed by incubation at 4C for 30 minutes with rotation. Chromatin was then pelleted by centrifugation at 25,000 x g for 30 minutes, resuspended in buffer AP (15 mM HEPES, 60 mM KCl, 15 mM NaCl, 5% sucrose supplemented with 0.5 mM spermine, 0.15 mM spermidine, 1X PIC, 0.4 mM PMSF, and 1 mM BME), and treated with 0.4 U/mL MNase at 37C for 30 minutes. Digested material was then pelleted, resuspended in 5 mM EDTA with 0.2 mM PMSF, and brought to 0.75 M NaCl by adding 5M NaCl dropwise. Nucleosomal material was then layered over 35 mL sucrose gradients (5-30% sucrose in 15 mM HEPES, 0.75 M NaCl, 1 mM EDTA, 0.5 mM PMSF, and 1 mM BME) and centrifuged in a SW-28 rotor at 26K rpm for 18 hours. Fractions were collected 1 mL at a time and those of suitable oligomer size (∼0.6-2kb, assessed by agarose gel) and purity (assessed by coomassie) were pooled, dialyzed overnight against 4 L of oligonucleosome storage buffer (20 mM Tris 8.0, 100 mM NaCl, 1 mM EDTA, 10% glycerol, 4 mM PMSF, and 1 mM DTT), then against 2 L of storage buffer for 3 hours, clarified by centrifugation, and quantified by nanodrop before flash-freezing in 5 µM aliquots.

### PRC2 purification for biochemical assays

Native PRC2 complexes were purified from HEK293 cells expressing FLAG-tagged EED, tagless EZH2, RBBP4 (for four-subunit PRC2), and the indicated tagless SUZ12 constructs. Nuclei were isolated by resuspending cells in hypotonic lysis buffer A (20 mM HEPES pH 7.9, 2 mM MgCl_2_, 10 mM KCl, 0.1 mM EDTA, 0.5 mM EGTA supplemented with 1 mM benzamidine, 0.8 mM PMSF, and 1 mM DTT) followed by douncing. Pelleted nuclei were resuspended in buffer AB (20 mM HEPES pH 7.9, 110 mM KCl, 2 mM MgCl2, 0.1 mM EDTA supplemented with 1X EDTA-free protease inhibitor cocktail (Pierce), 0.4 mM PMSF, and 1 mM DTT.

Nuclear extract was prepared by adding 1/10th volume of saturated ammonium sulfate followed by incubation at 4C for 30 minutes with rotation. Chromatin was pelleted by ultracentrifugation at 125,000×g for 90 min. Soluble extract was precipitated with a 60% ammonium sulfate cut by adding 0.3 g of solid ammonium sulfate per mL of extract over 5 minutes in a cold room with stirring. After 20 more minutes of stirring to allow for complete precipitation, precipitated extracts were pelleted by centrifugation at 20,000 x g for 30 minutes, flash-frozen, and stored at -80C overnight.

Extract pellets were resuspended in a suitable volume of binding buffer (20 mM HEPES 7.9, 300 mM KCl, 1 mM EDTA, 0.01% NP-40, 10% glycerol, 0.4 mM PMSF, 1 mM BME) and dialyzed against 2 L of binding buffer twice for 2 hours each. Dialyzed extracts were incubated with a suitable volume of M2 FLAG beads (Sigma) for 4 hours at 4C. Protein-bound beads were collected by centrifugation, resuspended in wash buffer WB-1000 (20 mM HEPES, 1 M KCl, 1 mM EDTA, 0.01 % NP-40, 10% glycerol, 0.4 mM PMSF, 1 mM BME), transferred to a gravity flow column, and then washed with 30 column volumes of WB-1000. Beads were lastly washed with 10 column volumes of WB-400 (identical to WB-1000 but with 400 mM KCl). Complexes were eluted in WB-400 supplemented with 0.3 mg/mL 3xFLAG peptide (Tufts University Peptide Core Facility). FLAG-purified material was then passed over a DEAE column and finally purified by cation exchange chromatography (mono S). All purified complexes were stored in 20 mM HEPES 7.9, 1 mM EDTA, 10% glycerol, 400 mM KCl, 2 mM BME, and 0.4 mM PMSF at a final concentration of 4 µM.

### Histone methyltransferase (HMT) assays

Typical HMT reactions were performed in a total volume of 20 µl containing HMT buffer (25 mM Tris 8.0, 2 mM MgSO_4_, 5 mM DTT, 0.4 mM PMSF), 1.42 µM ^3^H-SAM (0.5 µCi) (Perkin Elmer), 2.58 µM cold SAM, 200 nM oligonucleosomes, 20 nM PRC2, 40 µM H3K27me3 stimulator peptide, and 100 mM monovalent salt (typically 24 mM NaCl from nucleosome buffer, 76 mM KCl added). For RNA-containing reactions, 2 U of RNase inhibitor (NEB) was added. Reactions were typically carried out for 45 minutes at 30C; all time points were determined to be within a linear range in preliminary assays. Experiment-specific modifications are noted in figure legends. For scintillation, reactions were spotted on phosphocellulose filter paper, dried for 30 minutes, then washed 3x in 0.1 M NaHCO_3_ for 30 minutes each. Washed papers were dipped in acetone and dried fully by air. Scintillation counting was performed using a Tri-Carb 2910 TR liquid scintillation analyzer (Perkin Elmer). Counts were corrected for background and automethylation using reactions without substrate. For IC50 determinations, percent activity values were calculated by normalizing to reactions without nucleic acid inhibitors performed in the same experiment; graphed data points represent percent activity values from at least three independent experiments.

### Electrophoretic mobility shift assays (EMSAs)

Binding reactions were performed in a total volume of 10 µl containing 50 mM Tris pH 7.6, 200 mM KCl, 0.025% NP-40, 2 mM MgCl_2_, 0.1 mg/mL BSA, 10% glycerol, 0.4 mM PMSF, and 1 mM DTT. Reactions were incubated for 30 minutes at 30C to reach equilibrium. Reactions were then ran through a native 0.7% agarose gel buffered in 1X TBE for at 6 V/cm in a cold room for 45 minutes. Fluorescent signals were captured by imaging with a Typhoon 5. Total signal intensities for bound or unbound bands were quantified by measuring integrated density signals with ImageJ. Percent bound values from three independent experiments were plotted in GraphPad Prism 10 software and Kd values obtained by fitting the data to a four-parameter Hill curve with variable slope (specific binding).

### High-throughput chromosome conformation capture (Hi-C)

Cells were pelleted, washed with PBS, then crosslinked with 1% PFA in PBS for 10 minutes at room temperature with end-over-end rotation. Reactions were quenched by adding 200 mM glycine and incubating for 5 minutes at room temperature with rotation. Crosslinked cells were pelleted, washed with PBS, then flash-frozen and stored at -80C. DNA was then digested within intact, permeabilized nuclei using a 4-cutter restriction enzyme (DpnII). Resulting overhands were filled in, biotinylated, and blunt ends ligated. DNA was then sheared to a size of 300-500 bp by sonication (Covaris) with the following conditions: duty cycle 15, PIP 500, Cycles/Burst 200, 58 seconds. Biotinylated ligation junctions were then pulled down using streptavidin beads. Sequencing libraries were amplified from beads in 200 μl of PCR amplification mastermix (KAPA HiFi Hotstart ReadyMix, and KAPA Primer Mix), purified with AMPure XP beads, and sequenced for approximately 350M reads on an Illumina NovaSeq6000 S4 v1.5 PE150 platform.

### RNA-seq

RNA was extracted by resuspending cell pellets with 800 µl of TriZol reagent and nucleic acid was purified by addition of 160 µl of chloroform and centrifugation. RNA was isolated using a Quick-RNA Purification kit (Zymo Research) and genomic DNA was removed using Turbo DNase treatment for 15 min followed by an additional RNA column purification. Sequencing libraries were generated using NEBNext Ultra RNA Library Prep kit with Poly(A) mRNA isolation according to manufacturer instructions. RNA from two independent biological replicates were sequenced. Libraries were sequenced on an Illumina NovaSeq X Plus 1×50bp at approximately 20×10^6^ reads per sample.

### Clonal knockout generation

Cells were transiently transfected with pX459 gRNA plasmids using the GenJet transfection reagent. Following selection with puromycin, single cells were sorted into 96-well plates by FACS under sterile conditions. Clones were screened for knockout by PCR and verified by whole-cell immunoblot. All clonal knockouts were determined to be mycoplasma-negative and confirmed to have lost the transiently transfected CRISPR plasmid by reacquisition of puromycin sensitivity.

### Co-immunoprecipitation

Analytical co-IPs were carried out using extracts derived from 293F SUZ12-KO cells expressing FLAG-tagged SUZ12 constructs at physiologic levels. Cells were pelleted, washed with PBS, then resuspended in hypotonic lysis buffer A (20 mM HEPES pH 7.9, 10 mM KCl, 2 mM MgCl_2_, 0.1 mM EDTA, and 0.5 mM EGTA supplemented with 1 mM benzamidine, 0.8 mM PMSF, 1 mM DTT) followed by douncing. Nuclei were then pelleted and soluble nuclear proteins extracted by resuspending in buffer C (20 mM HEPES 7.9, 0.42 M NaCl, 2 mM MgCl_2_, 0.2 mM EDTA, 25% glycerol, 0.5 mM DTT, 0.5 mM PMSF, 1X protease inhibitor cocktail (Pierce)) and incubating for 30 minutes at 4C with rotation. Chromatin was pelleted by centrifugation at 21,000 x g for 30 minutes and supernatants (extracts) were diluted by adding 5 volumes of buffer D (20 mM HEPES, 180 mM KCl, 0.2 mM EDTA, 1 mM DTT, 1 mM PMSF). Anti-FLAG beads (M2, Sigma) were added and incubated at 4C for 4 hours with rotation. Protein-bound beads were collected by centrifugation, resuspended in wash buffer (20 mM HEPES 300 mM NaCl, 1 mM EDTA, 0.01% NP-40, 1 mM BME, 0.5 mM PMSF) and transferred to a micro-spin column. Beads were washed with 40 total column volumes before eluting in wash buffer with 0.3 mg/mL 3xFLAG peptide (Tufts University Peptide Core Facility). Eluates were normalized by FLAG levels as determined by immunoblot before assessing subunit co-IP efficiency.

## Acknowledgements

This research was supported by funding from the National Institutes of Health, including F30 NRSA fellowship F30CA257747 to T.J.R., R01CA266861 to P.W.L., P01CA196539 to P.W.L., N.J., and B.A.G., and P30CA014520 to the UW Carbone Cancer Center. Additional support was provided by an Innovator Award from Alex’s Lemonade Stand Foundation to P.W.L., grants from the Childhood Brain Tumor Foundation and the Rally Foundation for Childhood Cancer Research to P.W.L., and grants from the Sigrid Jusélius Foundation, the Emil Aaltonen Foundation, and the Otto A. Malm Foundation to J.K.L. P.W.L. is a Pew Scholar in the Biomedical Sciences. We thank Jim Keck for providing the G-quadruplex DNA oligonucleotide and Michael Erb for the dTAG reagents.

## Author Contributions

T.J.R. and P.W.L. conceived the study and wrote the manuscript. T.J.R. performed all experiments and analyses unless otherwise indicated. P.C. conducted all mouse experiments. A.B. analyzed Hi-C data. J.K.L. processed mass spectrometry samples. C.R. processed Hi-C samples and generated Hi-C sequencing libraries. A.Q.R. assisted in substrate preparation for biochemical assays. T.J.D. generated SUZ12-knockout 293F cells. B.A.G., C.L.K., N.J., Z.S.M., and P.W.L. supervised experiments.

---

**Figure S1.**
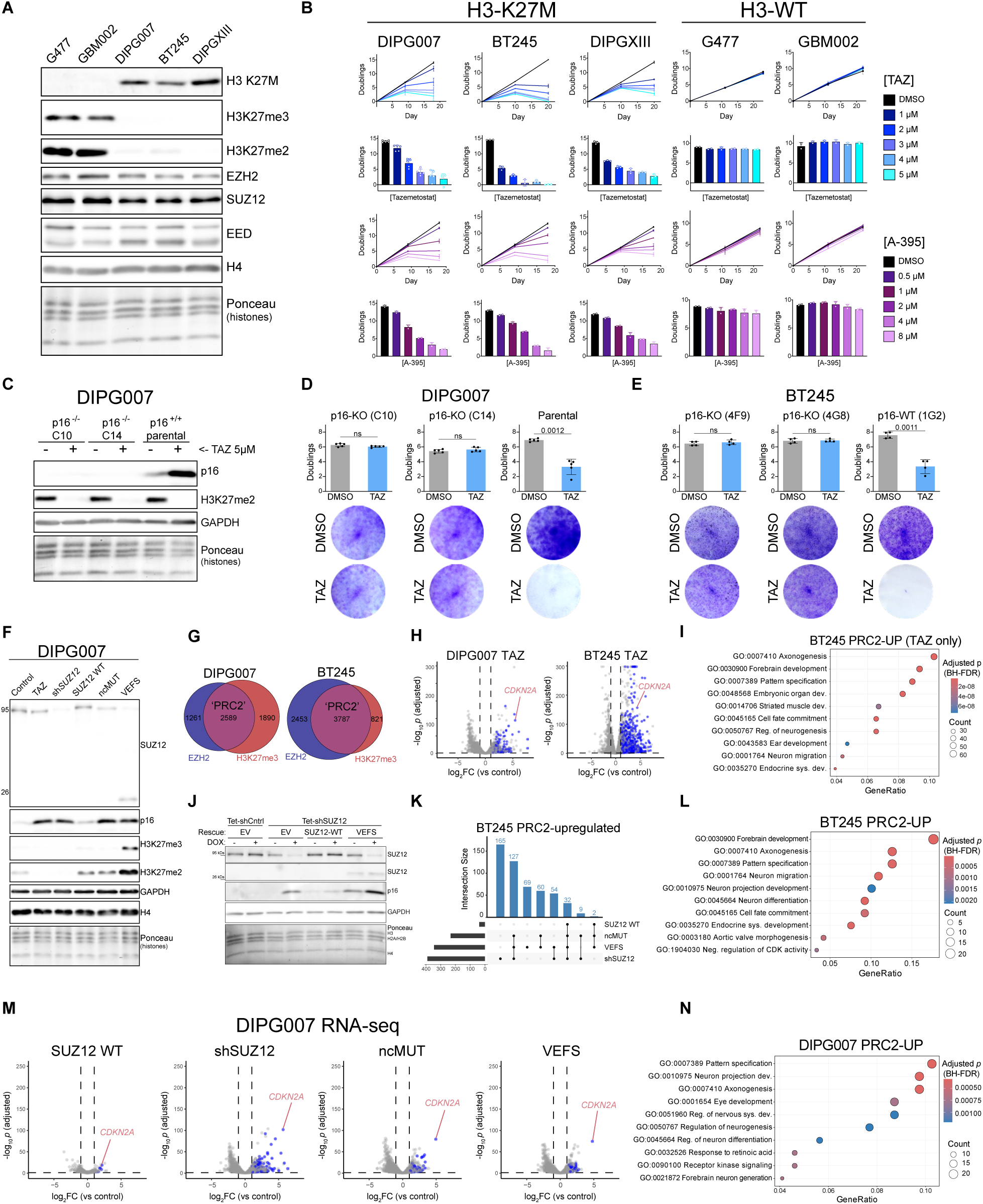
Supplementary Data Related to Figure 1. **(A)** Immunoblot analysis of whole-cell lysates from the indicated parental, patient-derived high-grade pediatric glioma cell lines utilized in this study. Antibodies used for each blot are indicated on the right; ponceau staining is included as an additional loading control. **(B)** In vitro proliferation assays evaluating sensitivity of each cell line to pharmacological inhibition of PRC2 enzymatic activity. Cells were treated with increasing concentrations of the EZH2 inhibitor Tazemetostat (TAZ, top panel) or the EED-binding allosteric inhibitor A-395 (bottom panel). For each line, cells were re-plated once as control (DMSO-treated) samples reached ∼80-90% confluency to avoid confounding effects of contact inhibition. Seeding densities at day 0 and at re-plating were equal across all conditions. Represented values are population doublings calculated from cell counts; XY plots represent each time point while the barplots represent cumulative doublings at the final time point. **(C)** Immunoblot analysis of whole-cell lysates from p16^INK4A^ knockout (p16 KO) cell lines. Shown are blots from two representative DIPG007 KO clones (C10 and C14) treated with or without Tazemetostat for 5 days alongside parental DIPG007 for comparison. Antibodies used for each blot are indicated on the right; ponceau staining is included as an additional loading control. **(D-E)** *In vitro* proliferation assays evaluating the response of p16 KO clones to Tazemetostat treatment over a 9-day period in DIPG007 **(D)** and BT245 **(E)** cell lines. For each DMG cell line, two KO clones are included (C10 and C14 for DIPG007; 4F9 and 4G8 for BT245) and p16^+/+^ controls for comparison (for DIPG007, bulk parental cells were used; for BT245, a clone (1G2) derived from the same process but found to be homozygous wild-type for p16 was used). Top panels show quantification of population doublings determined by direct cell counting. Bottom panels display representative images of crystal violet-stained monolayers from the same experimental conditions, providing a qualitative assessment of relative growth. Statistical significance was determined using two-tailed t-tests with Welch’s correction. “ns” denotes not significant (p > 0.05), indicating loss of sensitivity in p16 KO clones. **(F)** Immunoblot analysis of whole-cell lysates from DIPG007 cells with the indicated rescue conditions. Note that “TAZ” corresponds to “Control” transgenes and treatment with 5 µM Tazemetostat for 9 days. Lysates used for these blots were taken at the time of ChIP-seq and RNA-seq sample processing. Antibodies used for each blot are indicated on the right. Note that the SUZ12 antibody recognizes a region within the VEFS domain, thus recognizing all endogenous and transgenic SUZ12 protein in these lysates. Ponceau staining is included as an additional loading control. Integer values to the left indicate approximate molecular weight in kDa. **(G)** Euler diagrams showing the overlap of MACS2-called ChIP-seq peaks for EZH2 and H3K27me3 in the indicated untreated control cell lines. Peaks were called separately for each dataset, and the intersection between EZH2- and H3K27me3-enriched regions was designated as “residual PRC2 sites.” These overlapping peaks were used to define a consensus set of PRC2-bound loci for downstream gene expression analysis, including the annotation of PRC2-associated genes in RNA-seq datasets. **(H)** Volcano plots depicting transcriptomic changes in BT245 and DIPG007 cells following pharmacologic inhibition of PRC2 catalytic activity with Tazemetostat. Fold-change (log₂) is represented on the x-axis and statistical significance (−log₁₀ p-value) on the y-axis. Hashed lines denote significance thresholds (p = 0.05 and log₂ fold change of ±1). Transcripts significantly upregulated near PRC2 residual peaks, as defined in **(A)**, are highlighted in blue. The *CDKN2A* locus is specifically labeled in red to emphasize its derepression following PRC2 inhibition. **(I)** Gene ontology (GO) over-representation analysis of PRC2-associated transcripts upregulated in BT245 cells upon Tazemetostat exposure (blue dots in **(H)**). Displayed are the top 10 gene ontology (GO) terms. p-values shown are Benjamini-Hochberg corrected (BH-FDR). **(J)** Representative immunoblots of whole-cell lysates from DIPG007 cells validating the SUZ12 replacement approach. SUZ12 was detected with an antibody recognizing the VEFS domain to indicate expression of VEFS truncation relative to full-length endogenous and transgenic SUZ12 constructs. For rescue, cells were transduced at an MOI of 0.2 and lysates were harvested after 7 days of doxycycline-induced knockdown of endogenous SUZ12. Cropped SUZ12 blots show full-length and VEFS bands from the same membranes and exposures. Antibodies used for each blot are indicated on the right; ponceau staining is included as an additional loading control. **(K)** UpSet plots showing overlap of differentially upregulated transcripts from PRC2-associated genes in BT245 cells under the indicated experimental conditions. The 127-event union set was used for overrepresentation analysis in **(L)**. **(L)** Over-representation analysis of the union of upregulated PRC2-associated genes in BT245 cells from SUZ12 knockdown, ncMUT, and VEFS conditions. Displayed are the top 10 gene ontology (GO) terms. p-values shown are Benjamini-Hochberg corrected (BH-FDR). **(M)** Volcano plots of control-normalized gene expression changes in DIPG007 cells. Highlighted in blue are differentially upregulated PRC2-associated genes as described in **(A)**. Fold-change (log₂) is represented on the x-axis and statistical significance (−log₁₀ p-value) on the y-axis. Hashed lines denote significance thresholds (p = 0.05 and log₂ fold change of ±1). **(N)** Over-representation analysis of the union of upregulated PRC2-associated genes in DIPG007 (blue dots in **(M)**). Displayed are the top 10 gene ontology (GO) terms. p-values shown are Benjamini-Hochberg corrected (BH-FDR).

**Figure S2.**
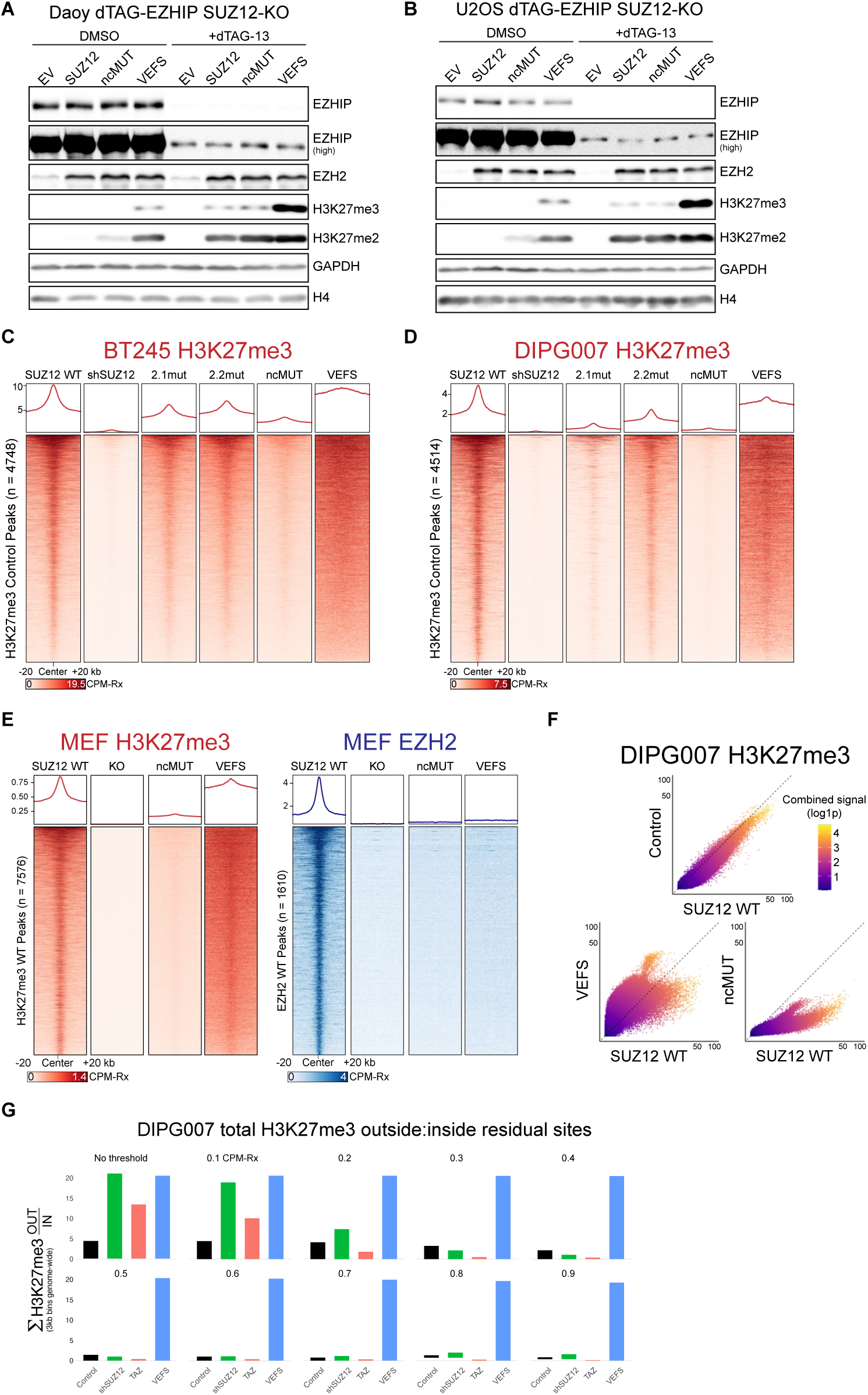
Supplementary Data Related to Figure 2. **(A-B)** Immunoblot analysis of whole-cell lysates from SUZ12-knockout, dTAG-EZHIP–expressing cell lines (DAOY medulloblastoma cells in A, U2OS osteosarcoma cells, in B) stably rescued with the indicated SUZ12 transgenes. Where noted, samples were treated with 500 nM dTAG-13 for 6 days to induce proteasomal degradation of the EZHIP fusion protein. **(B)** BT245 ChIP-Rx H3K27m3 signal at control peaks. The WT, shSUZ12, ncMUT, and VEFS data are identical to those in **Figure 2E** and are included here only for comparison to 2.1mut and 2.2mut samples. **(C)** BT245 ChIP-Rx H3K27m3 signal at control peaks. The WT, shSUZ12, ncMUT, and VEFS plots are identical to those in **Figure 2D** and are included here only for comparison to 2.1mut and 2.2mut samples. **(D)** Spike-in normalized ChIP-seq (ChIP-Rx) H3K27me3 signal in DIPG007 cells expressing the indicated SUZ12 rescue constructs. Heatmaps and associated metaplots are centered on peaks called in control cells. Metaplots display the average signal intensity across these regions in reference-normalized counts per million units (CPM-Rx). Samples were prepared and treated identically as the BT245 samples used to generate **Figure 2D**. **(E)** ChIP-Rx signal for H3K27me3 (left, red) and EZH2 (right, blue) in SUZ12 knockout MEFs expressing the indicated SUZ12 rescue constructs. Heatmaps and metaplots are centered on peaks called in SUZ12 WT rescue samples. **(F)** Scatterplots comparing genome-wide H3K27me3 ChIP-seq signal across 3 kb genomic bins. Each dot represents a bin, plotted by signal intensity in the two conditions being compared. The left panel compares the ncMUT and SUZ12 wild-type rescue; the right panel compares the VEFS and SUZ12 wild-type rescue. The color scale reflects the combined signal intensity across both samples, highlighting the distribution of differential enrichment and regions gaining or losing H3K27me3. **(G)** Bar plots in preliminary analyses in generating **Figure 2H**, showing the ratio of H3K27me3 ChIP-seq signal outside versus inside peak regions, calculated across a range of signal intensity thresholds. A range of thresholds was applied and visualized to identify low-signal noise that could significantly contribute to total signal intensity when taken in sum. Threshold values represent mean CPM-Rx per bin (for example, only bins with an average value of > 0.9 per million were included in the bottom right graph).

**Figure S3.**
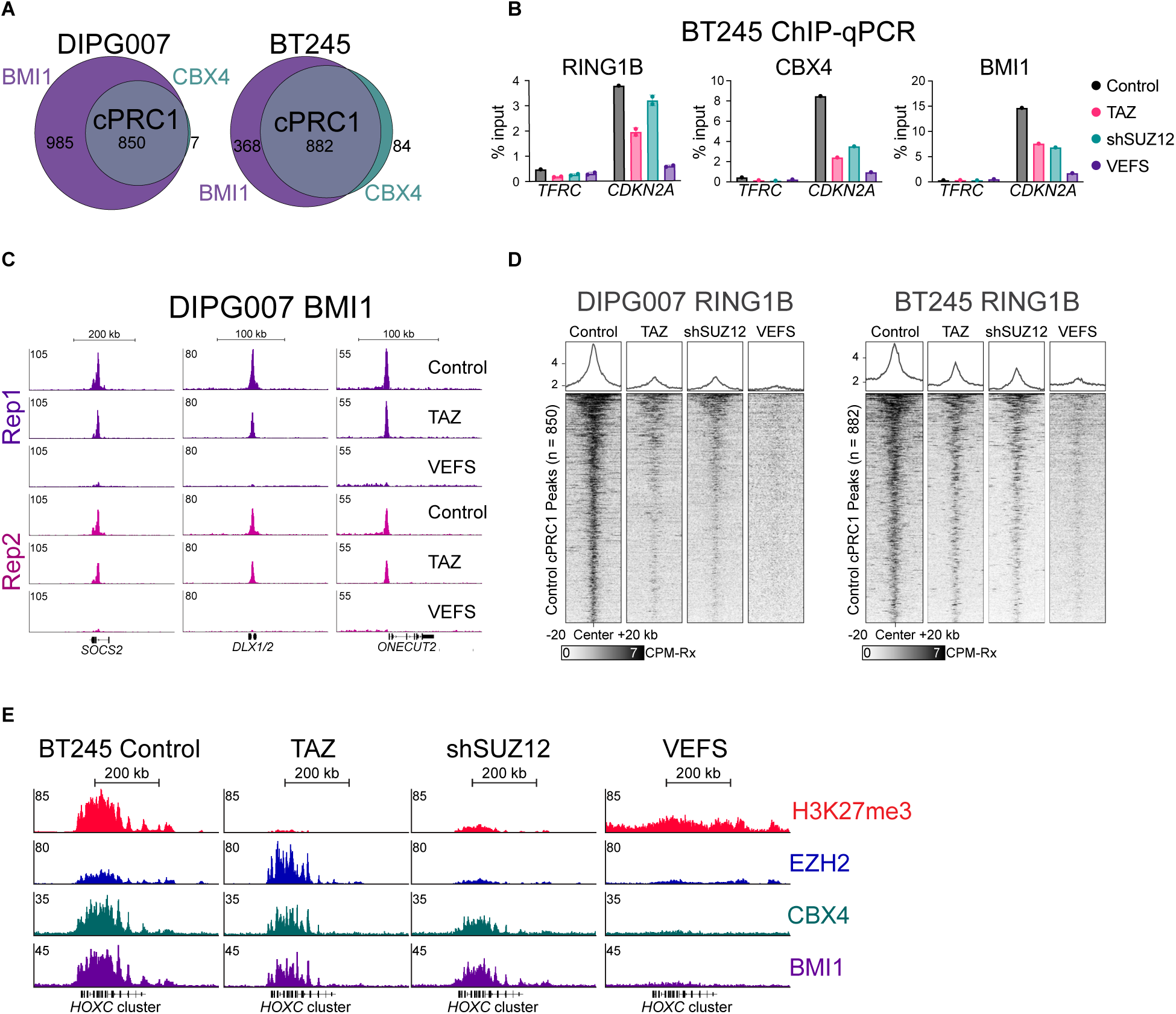
Supplementary Data Related to Figure 3 **(A)** Euler diagrams depicting the genomic overlap between CBX4 and BMI1 ChIP-seq peaks in control cells. The merged unions of these overlaps were used to define control cPRC1 peaks in Figure 3 analyses. **(B)** Quantification of BMI1, CBX4, and RING1B occupancy at the *CDKN2A* locus in BT245 cells using ChIP-qPCR. Enrichment values (percent input) were derived from technical replicates from independent experiments. A polycomb-negative site (TFRC) is included for comparison. **(C)** Genome browser tracks showing biological replicate ChIP-seq datasets for BMI1 in DIPG007 cells across the indicated experimental conditions. These replicate profiles confirm the reproducibility of BMI1 redistribution observed following loss of PRC2 catalysis (TAZ) or VEFS-driven hypermethylation. **(D)** ChIP-seq analysis of Ring1B, a catalytic core component of canonical PRC1, in multiple DMG cell lines under the indicated experimental conditions. Data are centered on cPRC1 peaks (as defined in panel A). **(E)** Genome browser views of ChIP-Rx signal for H3K27me3, EZH2, CBX4, and BMI1 in BT245 cells at the *HOXC* gene cluster, representative of densely-populated polycomb sites. See **Figure 3E** for similar visualization of *HOXD*. *HOXA* and *HOXB* clusters (not shown) exhibit similar patterns and levels of enrichment across all conditions

**Figure S4:**
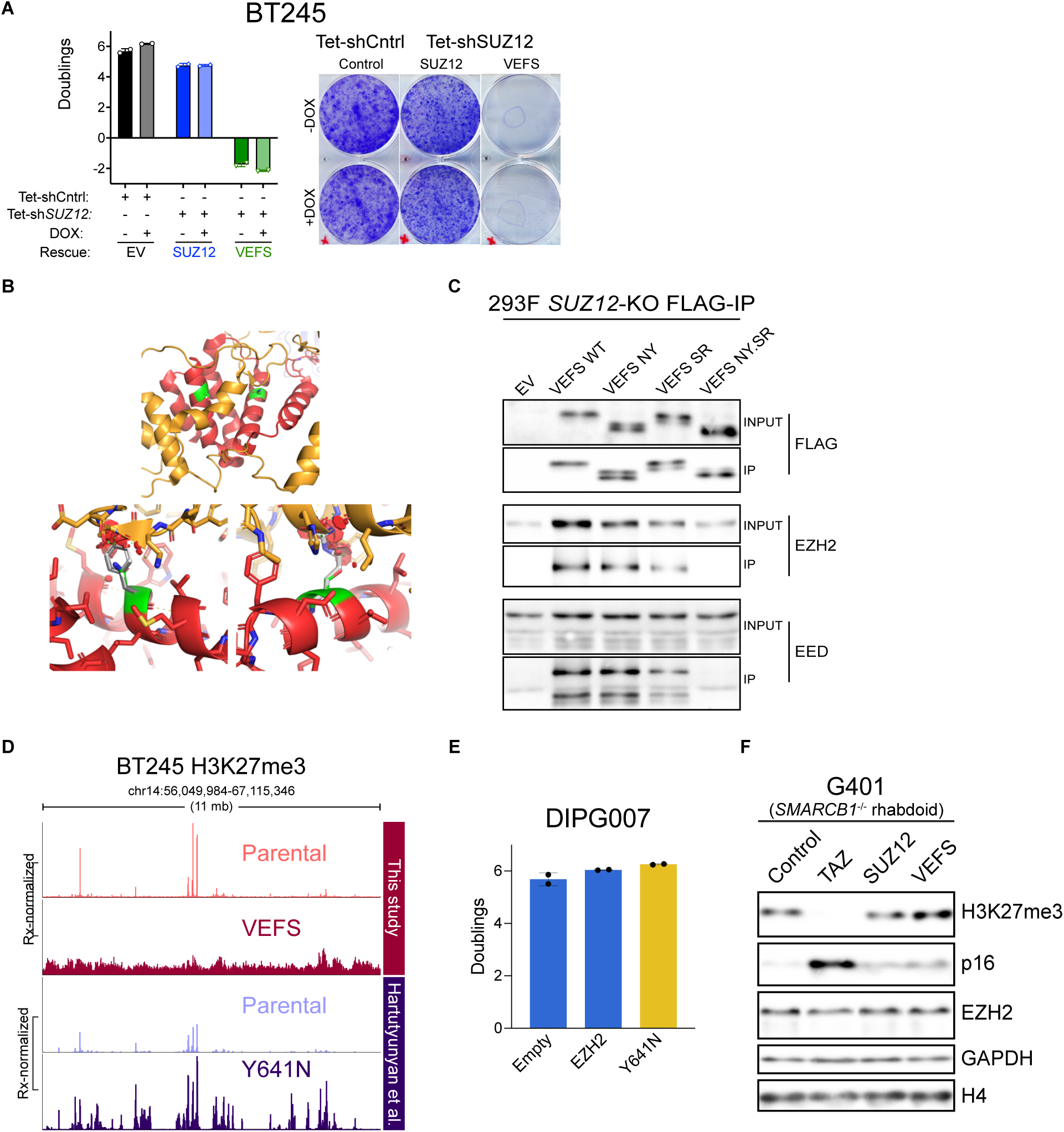
Supplementary Data Related to Figure 4 **(A)** Preliminary proliferation assay in BT245 cells stably transduced with the indicated SUZ12 rescue constructs at high multiplicity of infection (MOI), designed to ensure robust transgene expression. Proliferation was measured by direct cell counting (left) and visualized by crystal violet staining of monolayers (right). Both untreated and doxycycline-treated conditions are shown to evaluate the effect of shRNA-mediated knockdown of endogenous SUZ12. These data provide initial evidence that ectopic VEFS expression alone is sufficient to suppress cell growth, independent of endogenous SUZ12 depletion. **(B)** Structural modeling of the SUZ12 VEFS domain to evaluate the impact of engineered mutations that disrupt core PRC2 complex formation. PyMOL-rendered images based on the published SUZ12–EZH2 co-crystal structure (PDB 6C23) show SUZ12 (red) and EZH2 (orange). The wild-type interface residues N618 and S662 (top panel, green) are located at key contact points. Substitution of these residues with N618Y (bottom left) and S662R (bottom right) introduces predicted steric clashes (highlighted with red circles), likely disrupting physical interaction with EZH2 and destabilizing the core PRC2 complex. **(C)** Co-immunoprecipitation assays performed in SUZ12 knockout 293F cells to validate core-disrupting VEFS mutants. Cells were transduced with FLAG-tagged VEFS constructs (WT, N618Y, S662R, and N618Y/S662R double mutant). Immunoblots show that while individual mutations partially reduce co-IP of EZH2 and EED, the double mutation (referred to as NY.SR) is required to fully abrogate core complex formation. **(D)** Genome browser visualization of H3K27me3 ChIP-seq signal across an 11 Mb region on chromosome 14 in BT245 cells. The top two tracks represent data from this study under VEFS expression and control conditions; the bottom two tracks are from previously published datasets. All tracks are reference-normalized and scaled within each study. This visualization is meant to highlight qualitative differences in H3K27me3 distributions between SUZ12-VEFS and EZH2-Y641N despite globally similar levels in **Figure 4G**. **(E)** Proliferation assay in DIPG007 cells ectopically expressing either wild-type EZH2 or the hyperactive Y641N mutant. Cells were counted on day 9 and represent two biological replicates from one experiment. **(F)** Immunoblot analysis of whole-cell lysates from G401 rhabdoid tumor cells used in the proliferation assays presented in **Figure 4J**. Antibodies used for each blot are indicated on the right; H3K27me3 blots confirm VEFS-related hypermethylation without increased EZH2 expression. These cells are known to contain intact *CDKN2A* locus and repress p16 in a PRC2-dependent manner^104^. H4 and GAPDH are loading controls.

**Figure S5.**
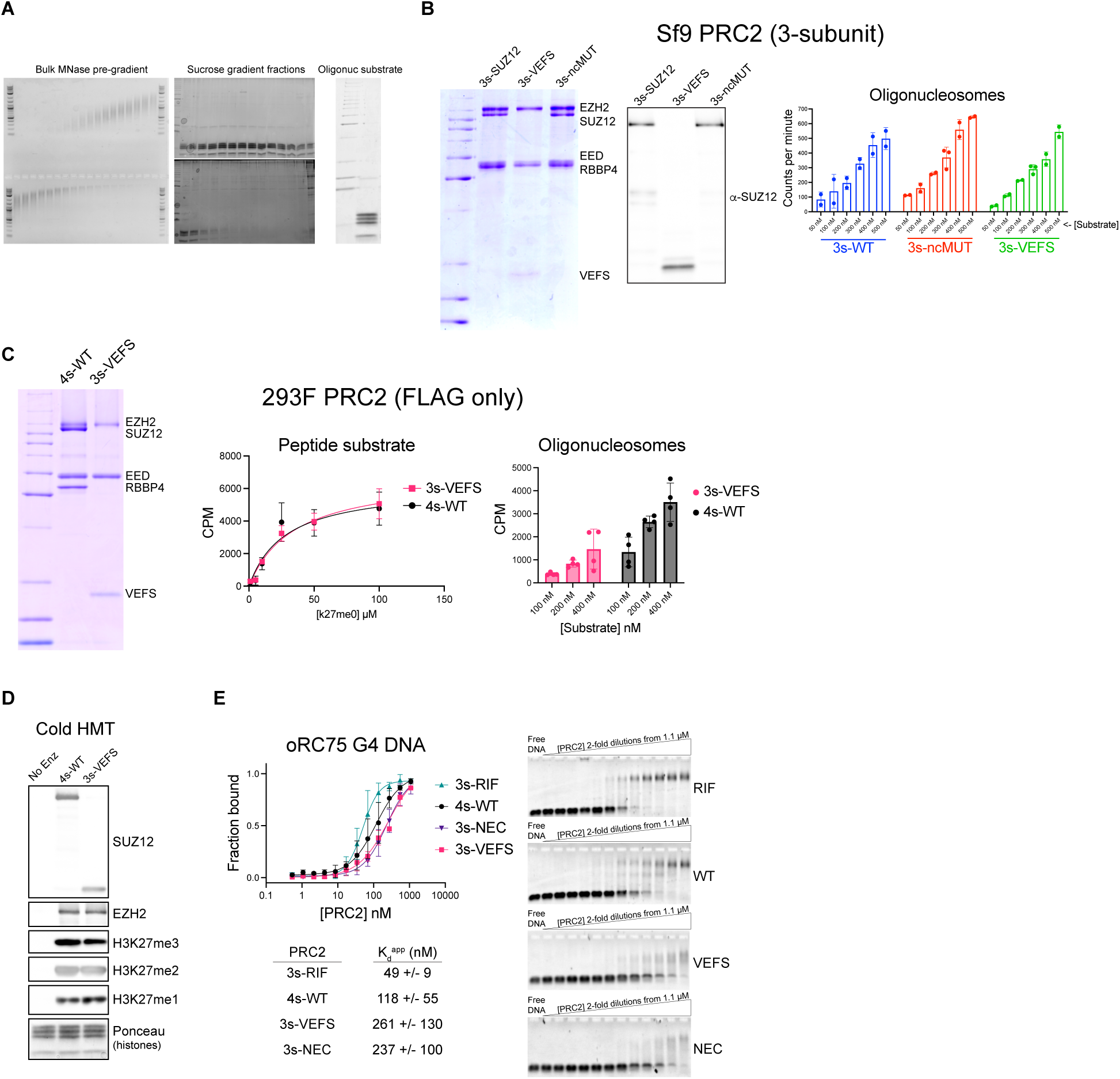
Supplementary Data Related to Figure 5 **(A)** Characterization of native oligonucleosome substrates used in HMT assays. Oligonucleosomes were purified from SUZ12-knockout 293F cells and confirmed to lack H3K27 methylation (data not shown). Left: ethidium bromide-stained agarose gel showing the integrity and size distribution of the extracted chromatin prior to fractionation. Right: Coomassie-stained SDS-PAGE of material collected from a sucrose gradient, confirming high-purity preparations of oligonucleosomes ranging from 4-12 nucleosomes in length. These pooled fractions were used as substrates in PRC2 activity assays throughout the study. **(B)** Comparison of HMT activity for Sf9-derived three-subunit PRC2 complexes reconstituted with different forms of SUZ12: full-length wild-type, VEFS truncation, or the full-length non-core mutant (ncMUT). Left: Coomassie-stained SDS-PAGE gel of purified complexes showing low overall protein yield and staining intensity for the VEFS-only complex. Middle: Western blot confirming equivalent molar loading of complexes despite differential staining in Coomassie. Right: preliminary HMT assays using 1 nM PRC2, 10 mM KCl, and nucleosome substrates, performed for 15 minutes. CPM, (scintillation) counts per minute corrected for background. **(C)** HMT activity of PRC2 complexes purified from pCAG 293F SUZ12-KO cells, as in the primary experiments shown in **Figure 5**. Complexes were used directly following FLAG affinity purification without further polishing. Left: Coomassie-stained gel showing protein composition of eluted complexes. Middle: activity on H3 tail peptides measured in reactions containing 10 nM PRC2 and 10 mM KCl for 45 minutes, with peptide concentrations titrated as indicated. Right: activity on native oligonucleosome substrates using 20 nM PRC2, 40 mM KCl, for 45 minutes. CPM, (scintillation) counts per minute corrected for background. **(D)** Immunoblot-based HMT assay using cold (non-radioactive) S-adenosylmethionine (SAM) to assess H3K27me levels produced by purified PRC2 complexes. Reactions were carried out to completion using 80 nM PRC2 and 1 µM native oligonucleosomes in 28 mM salt buffer for 3 hours. Reactions were stopped by mixing 16 µl of each reaction with 4 µl of 10X SDS/Laemmli buffer and boiling. SDS-PAGE and immunoblotting was carried out with antibodies indicated on the right. Ponceau staining is included as a loading control. **(E)** Binding of PRC2 complexes to a G-quadruplex–forming DNA probe assessed by EMSA. Left: quantification of DNA binding from three independent experiments. Right: representative agarose gels displaying shifted DNA–protein complexes.

